# Adaptive Milling: imaging feedback-driven automated fabrication of consistently thin cryo-lamellae

**DOI:** 10.64898/2026.09.20.752565

**Authors:** Thomas Glen, Thomas M. Fish, Elaine M. L. Ho, Patrick Cleeve, Luís M. A. Perdigão, Nadisha Gamage, Ravi Teja Ravi, James Boris Gilchrist, Michele C. Darrow, Mark Basham, Maud Dumoux, Casper Berger

## Abstract

Cryo-electron tomography of 100-200 nm lamellae prepared by focused ion beam milling enables structural analysis of macromolecules within cells, tissues, and small organisms, but lamella preparation remains time-consuming and limits throughput. Although automated workflows enable unsupervised fabrication, they commonly produce variable results because fixed, pre-calibrated parameters are applied without accounting for lamella-to-lamella variability. To overcome these limitations, we developed an Adaptive Milling approach that adapts milling parameters based on imaging feedback. We implemented this approach for the critical polishing stage as Adaptive Polishing, which uses machine-learning-based segmentation of scanning electron microscopy images to identify lamella features predictive of imminent collapse. These are used to periodically assess whether to continue or stop thinning during polishing. We performed a systematic comparison of the quality of lamellae prepared with Adaptive Polishing and conventional automation, and demonstrate that Adaptive Polishing produces thinner lamellae with greater consistency in thickness, and performs well for in situ cryo-electron tomography. Adaptive Polishing was tested on three different microscopes with minimal model retraining required, demonstrating transferability of the approach. We implemented Adaptive Polishing as an open-source plugin for fibsemOS together with machine-learning model training software and pre-trained models.

## Introduction

High-resolution cryo-electron tomography (cryo-ET) of thin cellular samples allows for the structural characterisation of macromolecules in their cellular context. The need for electron transparency typically limits cryo-ET to specimens no thicker than 300 nm. early studies therefore focused on naturally thin samples such as the periphery of eukaryotic cells (Medalia 2002) and small bacteria (Komeili et al. 2006). The development of sample thinning methods (McDowall et al. 1983), particularly the use of focused ion beams (FIB) to prepare 50-300 nm thick sections known as lamellae (Marko et al. 2007), has enabled in situ structural studies on a wide range of biological systems including the interior of mammalian cells (Mahamid et al. 2016), pathogens during intracellular infection (Böck et al. 2017; Wolff et al. 2020), tissue and small organisms (Schaffer et al. 2019). Combined advances in lamella preparation (Schaffer et al. 2017; Zachs et al. 2020), cryo-ET data acquisition (Eisenstein et al. 2023) and processing (Tegunov et al. 2021) have made it possible to obtain high-resolution (3-5 Å) in situ structures using cryo-ET (R. Kelley et al. 2026; Xing et al. 2023; Xue et al. 2022). To obtain these high resolutions, the preparation of large numbers of thin and stable lamellae is critical, with a thickness of 100-200 nm being a key determinant of data quality (Lucas and Grigorieff 2023; Tuijtel et al. 2024; R. Kelley et al. 2026). This requirement drives the need for high-throughput fabrication of consistently thin lamellae.

FIB lamellae are prepared in a series of milling steps. First the bulk of the material is removed at higher currents to maximise milling rates, followed by progressively lower currents, which provide finer control and produce smoother milling surfaces. In the final polishing step, lamellae are thinned down to a nominal thickness of 50-300 nm. Before any milling occurs, an organometallic platinum layer (Hayles et al. 2007) approximately 1-3 µm thick is deposited on the sample using a gas injection system (Wagner et al. 2020; Bisson et al. 2021; Berger, Ravelli, López-Iglesias, and Peters 2021). This sacrificial layer protects the lamella leading edge during milling and is often referred to as the GIS layer. Due to the intrinsic heterogeneity of biological samples and local variation in GIS deposition, optimal milling parameters typically differ between lamella sites.

Lamella fabrication was initially done manually, in a labour-intensive process in which an experienced operator can prepare around 10 lamellae per day (Schaffer et al. 2017; Medeiros et al. 2018; Wolff et al. 2019). During milling, operators often use scanning electron microscopy (SEM) and FIB imaging to monitor milling progress and adjust milling parameters according to the characteristics of each site. In particular, the removal of material is monitored, while tracking likely points of failure such as the local GIS layer thickness and the formation of cracks and holes. This is particularly important in the final polishing stages, where milling for several seconds too long can make the difference between a thin, smooth and intact lamella or the lamella being destroyed. More recently, automated milling approaches have been developed (Buckley et al. 2020; Zachs et al. 2020; Klumpe et al. 2021; Tacke et al. 2021) in which manually pre-calibrated parameters such as pattern sizes, milling duration and currents are applied to multiple milling sites. Unsupervised automated overnight milling allows for higher throughput of lamella fabrication of up to around 25 lamellae per day (R. Kelley et al. 2026). However, lamellae prepared with automated workflows often result in lamellae thicker than 200 nm (Supplementary Table 1), as pre-calibrated fixed parameters cannot account for site-to-site variation resulting from biological heterogeneity and variability in sample preparation. Optimising these protocols is also an iterative approach that generally requires experienced users.

Adaptive feedback microscopy (Hinderling et al. 2026) uses imaging feedback to automatically adapt microscope conditions to the sample. We reasoned this concept could be applied to lamella fabrication to adjust milling conditions for each lamella site, mimicking human decision-making.

In this study, we developed an Adaptive Milling approach for the final polishing step of lamella fabrication, termed Adaptive Polishing. This was done by training machinelearning segmentation models to identify lamella features predictive of imminent lamella breakage. We applied these models to reliably extract these features from periodically acquired SEM images during the critical polishing stage and developed rule-based decision-making to determine when to stop milling. We systematically compared lamella quality and milling success rate of Adaptive Polishing to traditional automation and found that Adaptive Polishing results in consistent preparation of significantly thinner lamellae on cells and obtained similar results when applied for preparing lamellae for in situ structural biology studies. We tested Adaptive Polishing on three different FIB/SEM microscopes using both plasma and gallium ion sources, and found that inclusion of only an additional 150 training images from a microscope that the segmentation model was not trained on is sufficient to greatly improve Adaptive Polishing decision-making. To make this method widely accessible, we implemented the Adaptive Polishing strategy as an open-source plugin for the vendor-agnostic FIB/SEM control software fibsemOS (Cleeve and Klumpe 2026) together with pre-trained models and software for model training.

## Results

### Adaptive Milling for the lamella polishing stage

To develop imaging feedback-driven Adaptive Milling for the fabrication of consistently thin (*<* 200 nm) lamellae, we focused on the final polishing step (Fig. 1a), the stage that determines the final lamella thickness and a likely point of failure during automation. This Adaptive Milling approach mimics how human operators decide when to stop polishing by periodically recording and analysing SEM images for lamella features predictive of imminent lamella collapse. Based on experiences in manual lamella fabrication, breakage of the GIS layer results in the formation of cracks (Lam and Villa 2021) and complete breakage of the lamella follows shortly after (Supplementary Fig. 1), regardless of precise lamella thickness. Monitoring GIS layer thickness during polishing can therefore help decide up to which point lamellae can still be safely thinned further. In some instances, small holes may form on the lamella surface while still protected by sufficient GIS, which restricts further thinning (Supplementary Fig. 1). We used both the minimum GIS thickness and the presence of cracks and holes as our main candidates for further testing.

**Figure 1.**
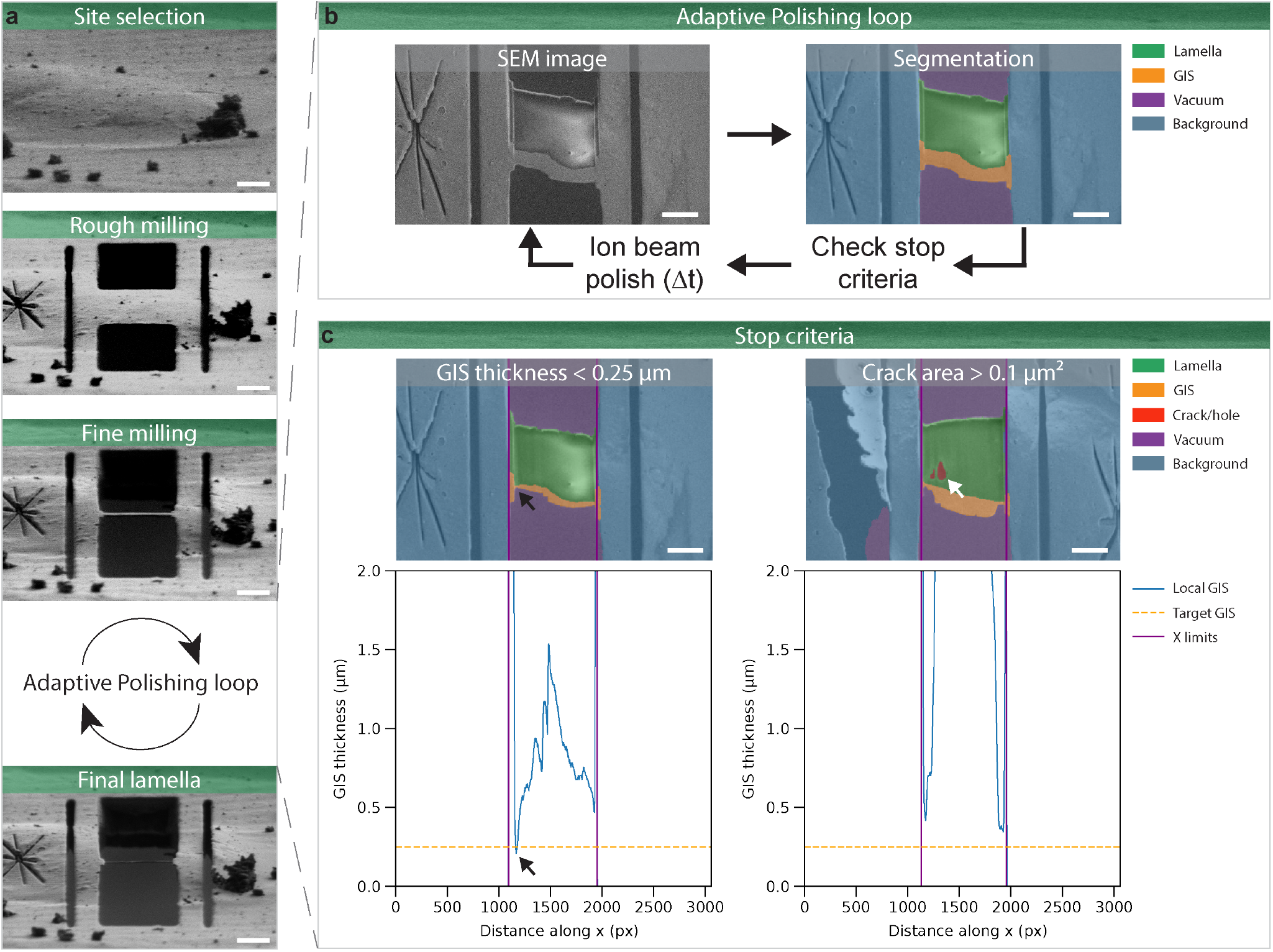
Adaptive Polishing uses SEM imaging feedback to determine polishing endpoints. (a) During automated milling, a series of milling patterns at progressively lower currents are used to remove all material above and below a section of the cell. Adaptive Polishing replaces the conventional automated polishing step where a fixed milling time is used. (b) Adaptive Polishing uses a loop consisting of ion beam polishing to thin down the lamella, SEM imaging and decision-making. The SEM image is segmented using a machine-learning segmentation model to identify the lamella surface, GIS layer, cracks/holes and two background labels (vacuum and other background). (c) These segmented images are analysed to determine the local GIS thickness along the width of the lamella (left) and the cracks/holes surface area (right). If the minimum GIS thickness is below a defined threshold (black arrows), or the cracks/holes surface area is above a threshold (white arrow), milling is stopped. If neither criteria are met, the lamella is further thinned using the ion beam (30 seconds in this example), and a new Adaptive Polishing cycle is started. Scale bars: 5 µm.

We developed an SEM-based Adaptive Milling approach for lamella polishing termed Adaptive Polishing (Fig. 1; see Methods section). Each polishing cycle consists of brief ion-beam milling followed by SEM imaging, machine-learning-based segmentation, feature measurement, and rule-based determination of whether polishing should continue. To measure lamella features, we trained a neural network (see Methods) to segment background, GIS layer, lamella surface, cracks and holes (segmented as one label named crack/hole), and vacuum (Fig. 1) using 2,873 SEM images of lamellae acquired during polishing using an Arctis FIB/SEM microscope, resulting in validation intersection over union (IoU; a commonly used metric for model accuracy) scores of over 0.6 for crack/hole (which are generally small and infrequent objects), and over 0.9 for all other labels (Supplementary Fig. 2).

To further improve the robustness of Adaptive Polishing, we implemented a routine that centres the lamella in the SEM field of view using beam shifts guided by the segmentation output (Supplementary Fig. 3). This compensates for offsets resulting from inaccuracies in the eucentric height or stage positioning and helps maintain the lamella and GIS layer within the imaged area. When using Adaptive Polishing for preparing lamellae it generally results in consistent segmentations and measurement of lamella features over time (Movie 1 and 2), supporting reliable rule-based decision-making. To support broad access to the method on different FIB/SEM systems, we implemented Adaptive Polishing as a plugin for fibsemOS (Cleeve and Klumpe 2026), which supports microscopes from multiple vendors. The Adaptive Polishing and other plugins are available in the open-source Adaptive Milling package with pre-trained models and training software (see Code & Data Availability).

### Adaptive Polishing consistently prepares thin lamellae

We determined the effectiveness of Adaptive Polishing by comparing lamella success rate, quality and milling throughput compared to conventional automation. We prepared lamellae with both methods on mouse embryonic stem cells (mESC) using an Arctis FIB/SEM microscope in three technical replicates. Each lamella was categorised as a “Success” (no or only small cracks or holes), “Partial” success (cracks or holes), “Failed” (e.g. destroyed during polishing, a crack over the full length of the lamella or large holes) or “Excluded” (e.g. broke during earlier milling steps or unable to reach stage position in TEM; see methods for full definitions). If possible, cryoEM tilt series were collected on all categories other than excluded. Both Adaptive Polishing and conventional automation reliably prepared lamellae, with 87% and 89% successful lamellae respectively (Fig. 2a; Supplementary Fig. 4). The thickness of the prepared lamellae was determined using cryo-ET, using a fixed acquisition pattern (Supplementary Figure 5) to avoid sampling biases and to account for the lamella wedge-like geometry (Schaffer et al. 2017; Berger et al. 2023, 2025; Tuijtel et al. 2024).

**Figure 2.**
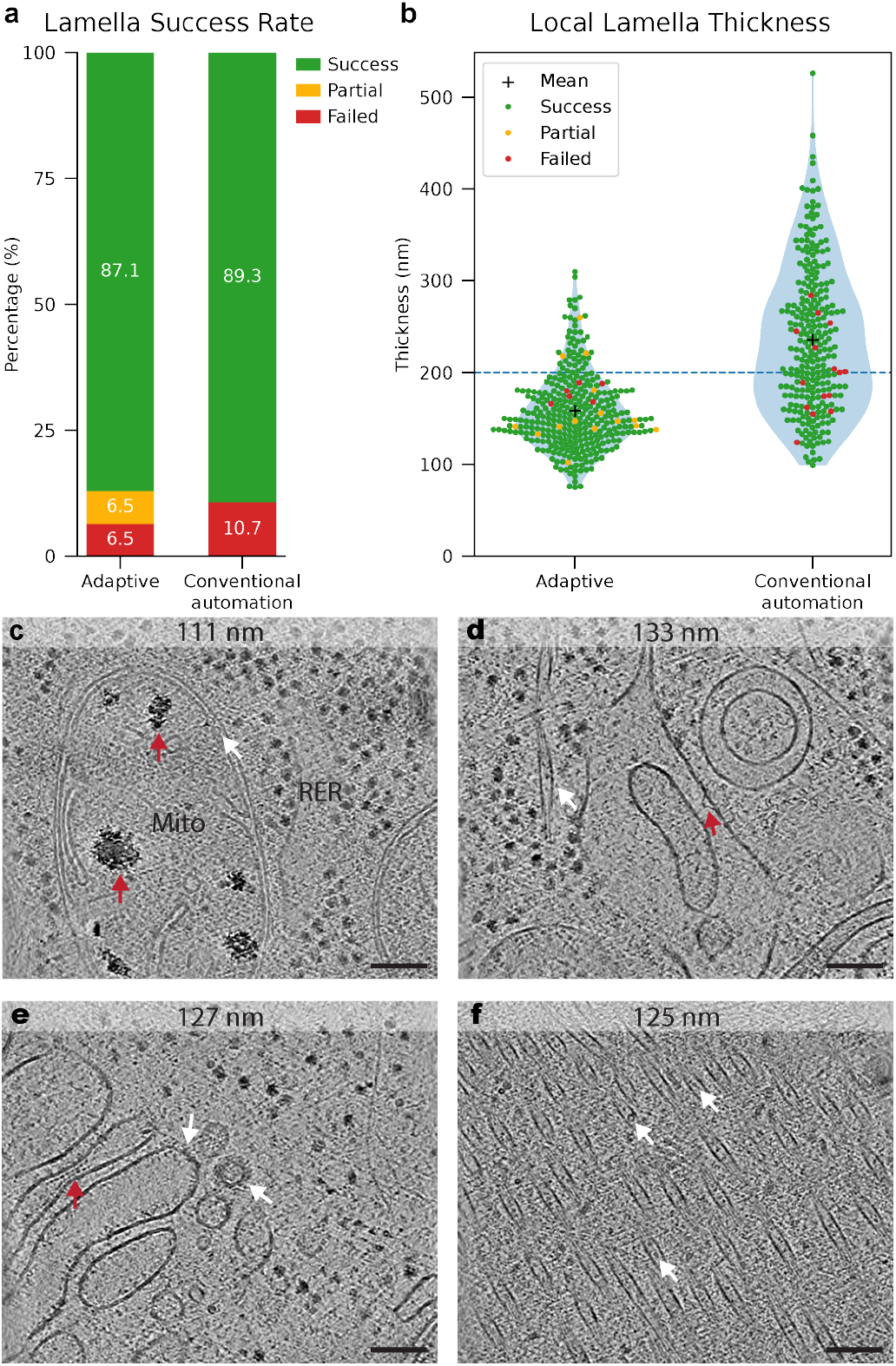
Experimental assessment of Adaptive Polishing performance. (a) Lamellae success rate for Adaptive Polishing (31 lamellae) and conventional automation (28 lamellae), showing the percentage of successful, partially successful and failed lamellae. (b) Local lamella thickness measured in tomograms for Adaptive Polishing (328 tomograms) and conventional automation (271 tomograms). Each tomogram is colour-coded according to the success condition of the lamella. The 200 nm thickness threshold is shown with the dashed horizontal blue line. Lamella thickness distributions were compared using a two-sided Welch’s t-test (t = -14.6, df = 394, P = 3.9 × 10^-39^). (c-f) Example tomograms prepared with Adaptive Polishing (c and d) or conventional automation with a fixed polishing time (e and f). The local lamella thickness is shown at the top. Scale bars: 100 nm. (c) Tomogram containing rough endoplasmic reticulum (RER) surrounded by ribosomes, as well as mitochondria (Mito). Inside one of the mitochondria, calcium-phosphate precipitates (red arrows) and macromolecular complexes such as Prohibitin (Rose et al. 2025) (white arrows) are also visible. (d) Tomogram containing a microtubule (white arrow) and an unidentified putative dome-shaped macromolecular complex (red arrow). (e) Tomogram containing the Golgi apparatus (Golgi), where vesicle budding and coated vesicles can be observed (white arrows). An array of putative macro-molecules also appears to constrict parts of the Golgi (red arrow), reminiscent of protein arrays observed previously at constrictions in Golgi cisternae in Chlamydomonas (Engel et al. 2015). (f) Tomogram containing a tight bundle of parallel microtubules. A large number of intraluminal densities can be observed within the microtubules (white arrows).

Adaptive Polishing produced significantly thinner lamellae that were more consistent in thickness compared to conventional automation (thickness±SD: 158±42 nm versus 235±78 nm; P *<* 0.001; Fig. 2b). Consequently, 86% of tomograms acquired on lamellae produced with Adaptive Polishing have local thickness below 200 nm, compared to 38% for conventional automation (Fig. 2b). A lamellae thickness below 200 nm has been demonstrated to be a critical factor for the resolution of structures obtained from cryo-ET data(Lucas and Grigorieff 2023; Tuijtel et al. 2024; R. Kelley et al. 2026). Tomograms acquired on thin lamellae with either Adaptive Polishing or conventional automation display a wide range of macromolecules and ultrastructural details (Fig. 2c-f; Movie 3-6). Next, we compared the total milling time of both methods. Adaptive Polishing only marginally increased the total preparation time per lamella (48 min 49 s versus 47 min 47 s), but reduced the milling time required per tomogram with a local thickness below 200 nm by more than half (5 min 5 s versus 12 min 12 s; Supplementary Fig. 6a).

To better understand Adaptive Polishing decision-making, we analysed the reasons it stopped milling. Most commonly, it stopped because the minimum GIS thickness threshold was reached (23/31 lamellae). The other lamellae stopped because either the cracks/holes surface area was above the threshold (5/31 lamellae) or because both the minimum GIS thickness and cracks/holes thresholds were exceeded (3/31 lamellae). The minimum GIS thickness of most lamellae was clustered around the predefined nominal 250 nm threshold (mean±SD; 237±232 nm; n = 31; Supplementary Fig. 6b) with much more varied median GIS thickness values (mean±SD; n = 31; 940±657 nm; Supplementary Fig. 6c). The per-lamella decision-making resulted in highly variable polishing durations (mean±SD 543±315 s; n=31; Supplementary Fig. 6d), with some sites stopped with less than 150 seconds of polishing time, and others after more than 900 seconds, illustrating the need for per-milling site modulation of the polishing time.

Together, these results show that Adaptive Polishing reproducibly prepares 160 nm thin lamellae, which are substantially thinner compared to conventional automation, and compares well to reported thickness values for both automated and manual polishing (Zachs et al. 2020; Berger, Ravelli, López-Iglesias, Kudryashev, et al. 2021; Klumpe et al. 2021; Li et al. 2023; Berger et al. 2023; Yang et al. 2023; Tuijtel et al. 2024; Schiøtz et al. 2024; Nguyen et al. 2024; Berger et al. 2025; R. Kelley et al. 2026) (Supplementary Table 1).

### Adaptive Polishing performs well for biological studies

We next evaluated Adaptive Polishing in a biological cryo-ET workflow using HeLa cells infected with Chlamydia trachomatis. In a 3-day milling session, 47 lamellae on 5 different grids were prepared using an Arctis FIB/SEM microscope, on which 714 tilt series were collected. We selected sites in the context of a biological study as opposed to the controlled site selection used in the comparison study on mESC cells. Adaptive Polishing yielded an 82.5% lamella success rate (Fig. 3a) and an average tomogram thickness of 150 ± 39 nm, with 89% of tomograms below 200 nm (Fig. 3b), using the same protocol as before. These values compare well to those obtained in mESCs and enabled high-quality visualization of intracellular ultrastructure and macromolecules (Fig. 3c and d). The lamellae are also among the thinnest datasets on cryogenic biological samples reported in the literature (Supplementary Table 1). These results demonstrate robust performance for Adaptive Polishing when applied for in situ cryo-ET studies.

**Figure 3.**
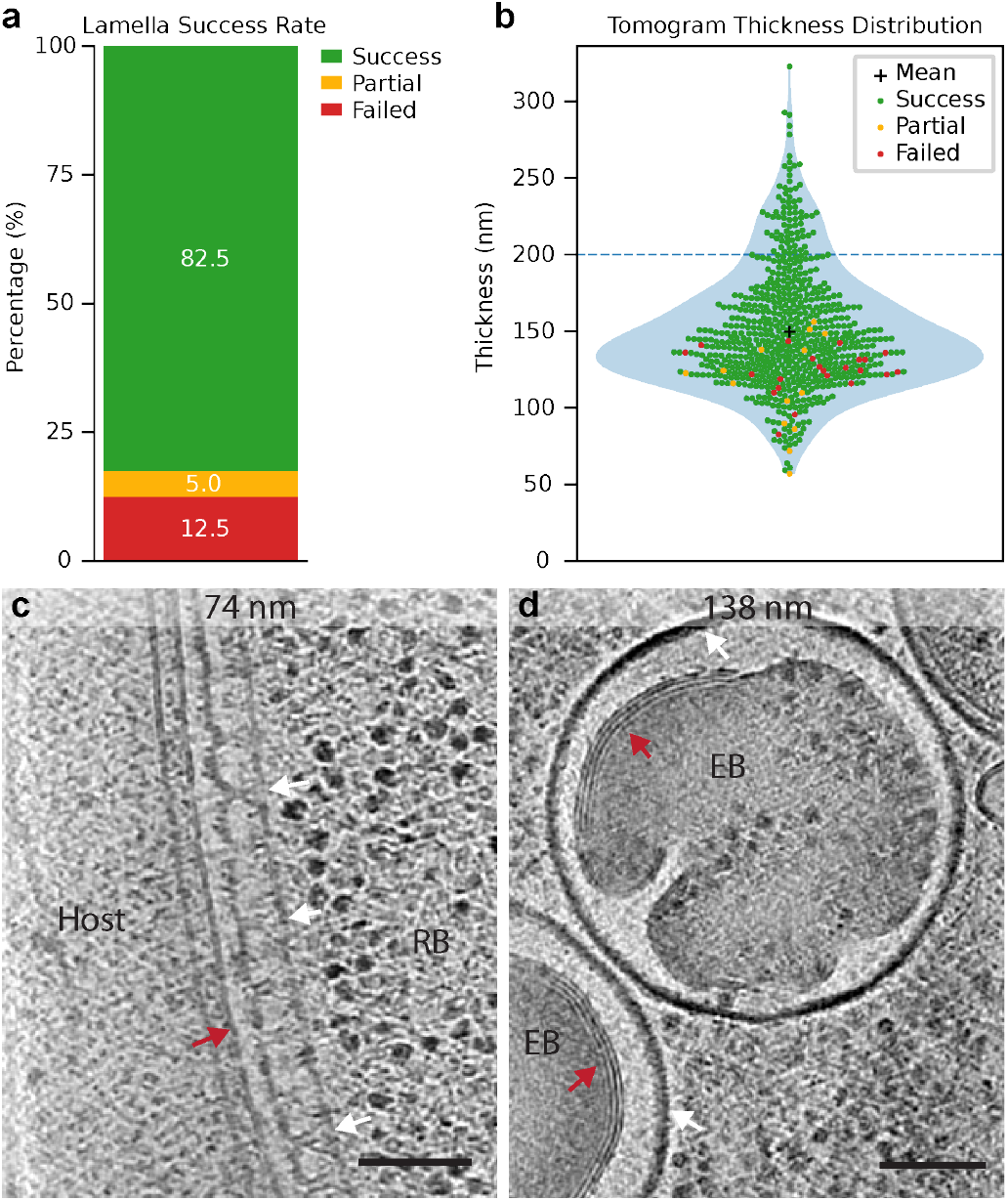
Adaptive Polishing on other cell types. (a) Lamellae success rate using Adaptive Polishing on Chlamydia trachomatis infected HeLa cells, showing the percentage of successful, partially successful and failed lamellae (40 lamellae total). (b) Tomogram thickness distribution (674 tomograms), where each tomogram is colour coded according to the success condition of the lamella. The 200 nm thickness threshold is shown with the dashed horizontal blue line. (c and d) Example tomograms prepared with Adaptive Polishing. Scale bars: 100 nm. (c) Reticulate body (RB) of C. trachomatis during intracellular infection of a HeLa cell (Host). An array of Type 3 secretion system complexes (white arrows) can be observed, facing the inclusion membrane (red arrow). (d) Elementary Bodies (EB) can be observed, with the characteristic thick outer membrane (white arrow). Unidentified stacks of high-contrast biological material (red arrows) can be observed along some regions of the inner membrane.

### Adaptive Polishing transfers well to other microscopes with minimal training data

We next assessed whether models trained on one microscope (Arctis) transferred to other FIB/SEM platforms. We prepared lamellae with Adaptive Polishing using an Aquilos 2 (using gallium) and a Helios Hydra (equipped with a different SEM column), using mESC and retinal pigment epithelial (RPE-1) cells respectively. While the model trained on only Arctis data still accurately segmented lamella features such as the vacuum-GIS and vacuum-lamella surface boundaries, indicating substantial transfer learning takes place, the GIS-lamellae surface boundary was often less reliably segmented and in some instances lipid droplets were incorrectly identified as small holes (Fig. 4a, first row). These mistakes in lamella segmentation often resulted in incorrect decision-making, such as early stopping (Supplementary Fig. 7).

To assess model performance after inclusion of microscope-specific data, we segmented additional SEM images (Aquilos 2: 145 SEM images from 7 lamellae; Helios Hydra: 161 SEM images from 19 lamellae) and used them together with the original data from the Arctis to train a new model (model name: “All”). Additionally, we trained 3 models by fine-tuning the “Arctis” model using combinations of Aquilos 2 and Helios Hydra data (model names: “Aquilos 2”, “Hydra” and “Aquilos 2 + Hydra”). As cracks and holes did not occur frequently in the acquired Helios Hydra dataset, we intentionally destroyed lamellae on some of the lamellae prepared with the Aquilos 2 by modifying stop condition threshold values (see Methods). We assessed model performance of the “Arctis” and the four new models by determining the IoU values for different labels using the held-back validation datasets from the three microscopes. We found that both training with all the data and transfer learning through fine-tuning with microscope-specific data resulted in more accurate label predictions for the Hydra and Aquilos 2 datasets compared to using the Arctis only model, while preserving performance on Arctis data (Fig. 4a).

**Figure 4.**
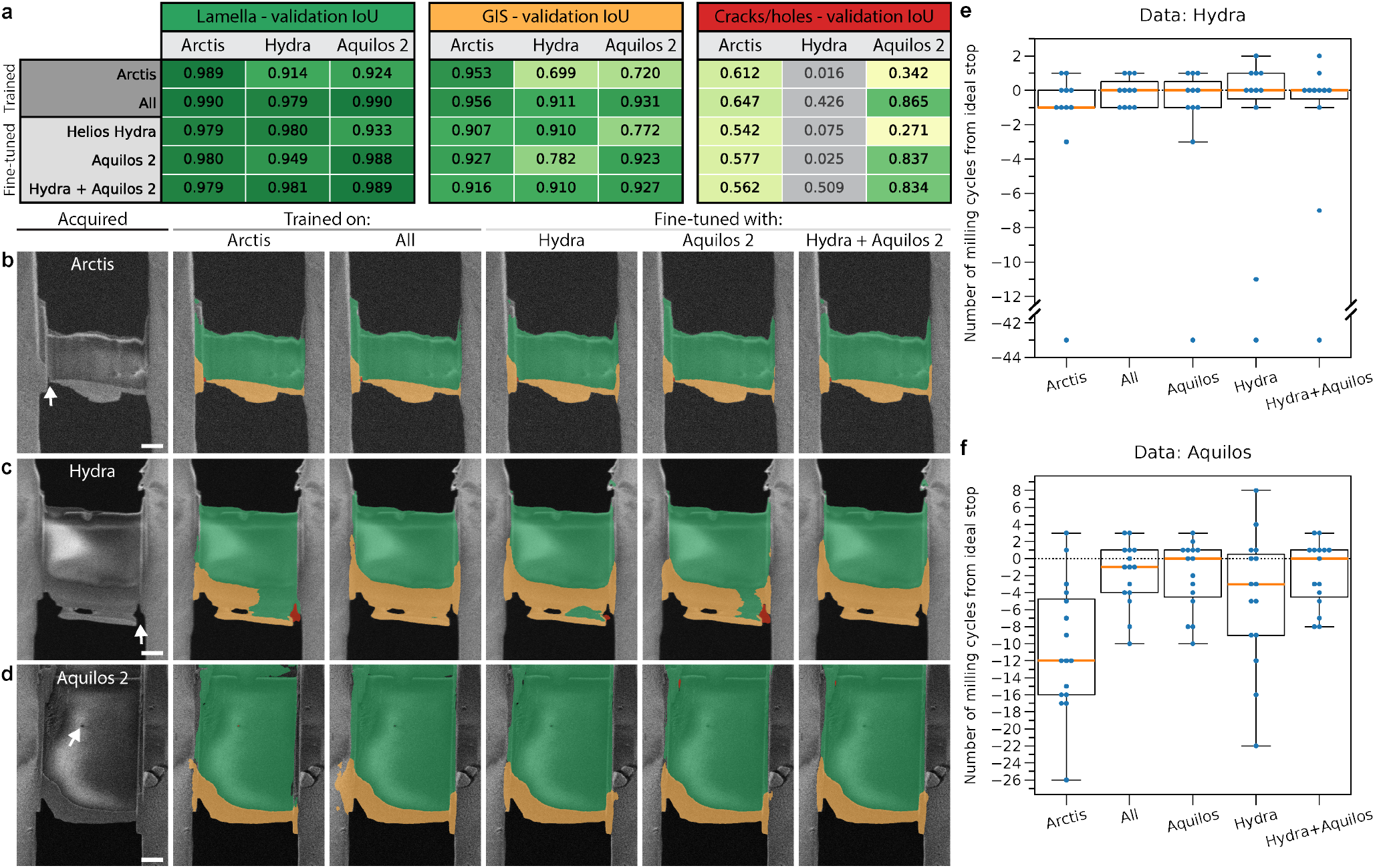
Improving model performance on other microscopes. (a) Validation IoU for the five segmentation models across the three microscope datasets for the labels of the lamella surface (green), GIS layer (orange) and cracks/holes (red). For the Hydra, performance in detecting cracks/holes could not be accurately assessed (shown in grey), as only a single SEM image contained a crack. (b-d) Representative segmentations without post-processing predicted by the five models on Arctis (b), Helios Hydra (c) and Aquilos 2 (d) datasets acquired after model training. Arrow in (b) indicates a small crack and arrows in (c) and (d) segmentation errors corrected by retraining or fine-tuning. Label colours: lamella surface (green), GIS layer (orange) and cracks/holes (red). Scale bar: 2 µm. (e and f) The box plots showing the difference in milling cycles between the cycle before a crack or hole forms and when Adaptive Polishing stops with each of the five different models for lamellae prepared with a Helios Hydra (e) or an Aquilos 2 (f). Median values (orange line) closer to zero indicate more accurate endpoint predictions.

To experimentally validate performance of these new models we prepared lamellae with the Hydra and Aquilos 2 using the “All” and “Aquilos 2” models respectively (Movies 7 and 8), using a second Adaptive Polishing step set to deliberately destroy the lamellae. We predicted labels with the 5 models on the SEM images and found similar segmentation improvements as previously seen on the validation data (Fig. 4b-d; Supplementary Fig. 8; Movies 9 and 10). To quantify the effect on Adaptive Polishing decision-making on this data for different models, we reran decision-making in silico for all 5 models and compared each model’s predicted stopping point to the cycle preceding crack or hole formation (Fig. 4e and f). Both fine-tuned and jointly trained models consistently stopped closer to this optimal endpoint compared to the “Arctis” model. During all the above experiments, we used the same imaging conditions on each microscope. We found that for the Aquilos 2, increasing the acceleration voltage to 2.5 to 5 kV greatly improves contrast between the GIS layer and the lamella (Supplementary Fig. 9) whereas for the Helios hydra, this was only the case for 5 kV (Supplementary Fig. 10). Changing imaging conditions such as the acceleration voltage can thus be used to increase contrast between the segmented lamella components, impacting model performance.

These results show that, incorporating approximately 150 microscope-specific SEM images during model training or fine-tuning is sufficient to transfer Adaptive Polishing to other microscopes. We also describe a strategy to deliberately destroy lamellae in order to rapidly collect SEM images containing cracks and/or holes for model training. To accelerate adaptation to additional microscopes, we provide pretrained models together with software for model training (Code and Data Availability).

## Discussions

In this study we describe Adaptive Milling for image-feedback-driven automated lamella fabrication. We implemented Adaptive Milling for the critical polishing step, and demonstrated its ability to consistently prepare 150-nm-thin lamellae. The development of Adaptive Milling methods accelerates the automation of in situ structural biology enabling larger datasets of higher resolutions, essential to make visual proteomics (Nickell et al. 2006) a practical reality.

The lamella thicknesses obtained with Adaptive Polishing were routinely among the thinner end of values reported in other studies (Supplementary Table 1). To avoid biasing the thickness measurements towards thinner regions, we acquired tilt-series in a fixed pattern on all lamellae where possible (Supplementary Fig. 5) and only excluded tomograms during processing if the thickness could not be reliably measured. In most other studies, this is typically not done, which may bias reported values towards lower thicknesses. It is also important to note that most studies do not report any metrics on lamella thickness, success rate or throughput.

While we show that the time-cost for Adaptive Polish is minimal (Supplementary Fig. 6a), the imaging overhead could be further reduced by only acquiring SEM images with a field of view containing the lamella and nearby context, and by dynamically adjusting the cycle length as a function of the risk of breakage. The latter can currently already be done by adding additional Adaptive Polishing steps before the final one in the protocol with safer thresholds and longer cycle lengths. For quantifying milling throughput, it is also important to consider how consistently thin lamellae can be prepared, as any time spent preparing thick lamellae is typically not productive towards the goal of acquiring high-quality cryo-ET data.

The observed transfer learning between microscopes suggests that a general model trained with data from diverse microscopes, milling applications and imaging conditions may enable robust Adaptive Polishing decision-making for most microscopes and use-cases. Decision-making performance for specific use-cases and instruments can still be further improved using model fine-tuning, where Adaptive Polishing can be used to rapidly capture training data of broken lamellae, as the most informative SEM images for model training are at the decision boundary rather than early in the polishing step or successful lamellae. It is also important to note that imaging conditions, such as acceleration voltage, can be changed to increase contrast between GIS and the cellular material to improve model performance on a given microscope (Supplementary Fig. 9 and 10). While Images of lamellae in different states are commonly collected during both manual and automated workflows, they are commonly not shared. General model training for Adaptive Milling implementations would benefit if this data was made available in public data repositories. In this study we focused on cells on grids, but we do not foresee fundamental issues in applying Adaptive Polishing to waffle-style(K. Kelley et al. 2022; Berger et al. 2025) and lift-out lamellae (Schiøtz et al. 2024; Nguyen et al. 2024; Glynn et al. 2025).

It is important to note that Adaptive Polishing prepares thin lamellae by thinning them down for as long as possible by detecting when further thinning will likely destroy the lamella, rather than measuring lamella thickness and using that as an endpoint. To obtain even thinner lamellae with Adaptive Polishing, milling could be controlled locally based on imaging feedback, similar to how a human operator may use exclusion zones to avoid milling areas at risk. While it is clearly established that thicker lamellae *>* 200 nm are detrimental for the resolution of obtained STA structures (Lucas and Grigorieff 2023; Tuijtel et al. 2024; R. Kelley et al. 2026), sample thickness affects many parameters, such as contrast, relative proportion of lamella surface damaged during FIB milling, lamella stability, the size of the continuous volume and the number of particles present in this volume per tomogram. Optimal sample thickness likely depends on the biological questions and the macromolecule being studies, where for example 100 nm thin lamellae may be preferred for smaller macromolecules such as nucleosomes, which are hard to reliably detect in thicker samples (Hou et al. 2023). An image-feedback mechanism for lamella thickness during milling would allow for the preparation of lamella with a specific thickness and provide quality assessment prior to cryo-ET data acquisition. This may be possible using a calibrated contrast in SEM images (Conlan et al. 2019; Schaffer et al. 2017), STEM techniques(Skoupý and Boltje 2023) or cryo-fluorescence imaging (Boltje et al. 2025) and has already been demonstrated using the backscattered electron signal on perovskite materials at room temperature (Tsurusawa et al. 2024).

In our current implementation, Adaptive Milling is only applied in the final polishing step. Applying imaging feedback-driven Adaptive Milling in earlier milling steps could bring substantial benefits. Current pre-calibrated protocols often use generous milling times to ensure all material is removed for every lamella site, sacrificing throughput for higher success rates. Manual calibration of protocols is also an iterative and time-consuming process that needs to be redone for different samples and generally requires experienced users. Robust Adaptive Milling for all steps could enable substantially higher throughput, improve reproducibility across different samples without the need for user parameter tuning and democratise access to FIB-milling by lowering training barriers. When Adaptive Milling for all stages is combined with automated lamella site selection (Klumpe et al. 2023), fully autonomous FIB milling could become a practical reality.

## Materials and Methods

### Proof of concept development

The concept of SEM feedback-driven automated milling was first prototyped using Python scripts with Autoscript 4 (Thermo Fisher Scientific) to interface with the FIB/SEM microscope. Lamella sites were prepared to a thickness ready for polishing using AutoTEM (Thermo Fischer Scientific), followed by manual pattern placement. As an initial proof of concept, the GIS thickness was analysed on automatically acquired SEM images using a manual line measurement and entry of the GIS thickness, and polishing was then conducted step by step, until the measured GIS thickness was reduced below a set threshold value. The GIS thickness measurements were subsequently automated using classical image segmentation methods (median filtering and Otsu thresholding (Otsu 1979)). However, the robustness of this methodology was not high enough to allow for reliable automation, so a more general method using machine-learning methods trained on the data collected previously was adapted. This approach allowed for full automation of the Adaptive polishing cycle until stopping criteria were met.

### Adaptive Polishing algorithm

The ‘AdaptivePolishing’ milling strategy is a plugin within the freely available software fibsemOS that operates as a loop, assessing the lamella via machine-learning segmentation of the SEM image and milling until one of the user-configured threshold values are hit. For each lamella, the following steps are excecuted:

1. Load the segmentation model (see Creation of the Segmentation Models).
2. Centre the SEM field of view on the lamella (see Lamella Centring).
3. Run the milling cycle until a stop criterion is met:
  a. Acquire FIB and SEM images.
  b. Run the segmentation model on the SEM image and clean up the result (see Segmentation Post-Processing).
  c. Run analysis on the resulting segmentation and save the measurements as metadata.
  d. Determine whether milling should continue by comparing the measurements to the threshold values.
  e. If the checks passed, mill the lamella.
  f. Save the segmentations, both the original and the cleaned version.
  g. Create and save plots summarising the individual cycle, along with the segmentations (see Quantification and Validation).
4. Save metadata and plots summarising the all milling cycles (see Quantification and Validation).

#### Lamella Centring (Step 2)

An SEM image is acquired and segmented, which is then used to define the lamella edges, from which the lamella centre can be derived as described in Segmentation Post-Processing. The lamella centre is then used to determine the beam shift required to centre the lamella in the field of view (Supplementary Fig. 3). This is done once per lamella to ensure that the analysis done on the lamella is not affected by the lamella being partially cropped.

#### Segmentation Post-Processing (Step 3b)

The post-processing of the segmentation is handled in two steps:

1. Cleaning up the segmentation labels.
2. Defining the lamella edges.

The segmentation is cleaned up by finding the largest single area that is made up of lamella, GIS, and crack/hole labels. This is done by masking all the areas labelled as lamella, GIS, or crack/hole, then finding the largest single object made up of these components. GIS and crack/hole labels that are not connected to this object can be considered invalid, so crack/hole labels found elsewhere are reclassified as vacuum, and other GIS and lamella labelled areas are reclassified as background. After this process, a prediction is returned that contains only one lamella and the attached GIS and cracks/holes, with the rest being classified as vacuum or background.

Using the mask of the largest single lamella-GIS-crack/hole object, the edge coordinates of the lamella are detected using the mask of the largest single lamella-GIS-crack/hole object. In this object, the lamella edges are defined as the 10th and 90th percentiles along the x axis. These percentiles were chosen via trial and error to ensure that potential segmentation inconsistencies do not significantly affect the edge definition without significantly cropping the lamella. This is used for an initial centring of the lamella via beam shift as well as defining the X-axis limits that are used for further processing.

#### Measurement of Stop Criteria (Step 3d)

GIS thickness is measured via the following steps:

1. The cleaned segmentation is cropped to the X-axis limits and from the top of the lamella, as defined by the post-processing, to ensure that only the area below the top lamella edge is considered.
2. Masks of the GIS and background are combined as the segmentation can be inconsistent at the edges. This is because the distinction between the GIS layer at the leading edge of the lamella and GIS-covered background is purely contextual.
3. There often is some background below the vacuum at the bottom of the image, which should be ignored, so this is removed by:
  a. Masking both the vacuum and crack/hole labels that are below the lamella.
  b. Finding the coordinates across the top of this vacuum and crack/hole mask.
  c. Using these coordinates to remove any background that is below these points.
4. The mask resulting from the previous steps is then padded to the full size of the segmentation before being resized to match the image dimensions. Bicubic resizing is used for this as it better considers morphology than a linear interpolation of the GIS thickness.
5. The output from measurement step 4 is then summed across the Y axis and multiplied by the image pixel size to find the physical thickness of the GIS layer along the X axis.

To determine the size of cracks and holes, the total surface area of the cracks/holes label is summed in the post-processed segmentation and converted to physical units via the pixel size. Note that all measurements are done as they appear in the SEM image, without considering the angle at which the SEM and the FIB are oriented relative to each other (52° in the microscopes used in this study).

#### Quantification and Validation (Step 3g and 4)

The milling cycle summary plot from Step 3g contains both the FIB and SEM images, along with the segmentations and a plot showing GIS thickness measurements (see Movie 1 and 2). These plots allow users to quickly inspect the end results of any automated milling that has been conducted, assess for any segmentation issues, and help the user determine whether the thresholds would benefit from an adjustment.

Results for each cycle, alongside other runtime metadata, are saved to a JSON file in Step 4 and can be used for more detailed analysis if necessary. Additionally, a plot summarising the GIS thickness measurements over time is created by Step 4, offering additional insight.

### Development and training of segmentation models

Cryo-SEM images were acquired during early script-based and later fully automated Adaptive Polishing on mouse embryonic stem cells (mESC) and hTERT RPE-1 Tet-On:Plk4 GFP-hCent2 cells. 8-bit or 16-bit greyscale images were recorded at 3072 × 2048 pixels, and in some cases multiple images of the same lamella state were taken under different imaging conditions (e.g. dwell time, defocus, acceleration voltage, and magnification). Images selected for annotation prioritised challenging cases where earlier model iterations performed poorly or where lamellae contained cracks or holes.

The first 500 images were manually annotated into five classes—background, GIS layer, lamella surface, cracks/holes, and vacuum. Vacuum regions were partially pre-segmented using the Segment Anything model (Kirillov et al. 2023) and refined in the openFIBSEM labelling UI (Cleeve et al. 2023) in Napari (Sofroniew et al. 2026) or labelled entirely manually. Subsequent annotations were created by manually correcting predictions from the best models at the time. All the training data and labels used in this study have been made available (see Data Availability). Segmentation training parameters, image preprocessing, and augmentations were iteratively improved during development, using the validation IoU for the lamella, GIS and cracks/holes labels as the main success criteria. The IoU is defined as the area of overlap between the prediction and expert annotation divided by the area of their union. Because IoU is normalized by object size, a discrepancy of only a few pixels has a much larger proportional effect for small objects than for large ones. Consequently, labels representing small and infrequent structures, such as cracks and holes, typically exhibit lower IoU values despite visually accurate segmentation. Current models were trained using a Feature Pyramid Network (FPN) with an EfficientNet-B6 encoder pretrained on ImageNet and implemented using the Segmentation Models library Pytorch(Iakubovskii 2019). Learning rate was regulated with AdamW optimiser (initial learning rate: 1 x 105; maximum learning rate: 1 x 103) with the encoder frozen for the first 25 epochs before unfreezing.

A compound loss function combining weighted cross-entropy (50%) and weighted multiclass Dice loss (50%) with class weighting, to mitigate class imbalances was used. The following class weights were used: Background (1.0), GIS (4.0), Lamella (3.0) Crack/hole (6.0) and Vacuum (2.0).

Training images were probabilistically augmented using random resized crop and/or gaussian blur, followed by normalisation (mean ±3 standard deviations to between -1 and 1), resizing to 1536 x 1024 pixels and converting to RGB. The probabilistic augmentation steps were skipped for validation images.

In total, 2,873 unique images (corresponding to 2,336 unique labels, as some lamellae timepoints were images with a few different imaging conditions that did not need to be individually annotated) were used to train and evaluate (15% random split) the original “Arctis” model in Adaptive Milling experiments. For training the “all” model and the three transfer learning models, additional proofread pseudo-labels were created for the Aquilos 2 (145 SEM images from 7 lamellae) and the Helios Hydra (161 SEM images from 19 lamellae). The “Arctis” and “all” models were trained for 100 epochs, with epochs 73 and 86 respectively chosen for subsequent experiments through standard overfitting approaches.

For model fine-tuning, the “Arctis” model was used as the initial model and retrained without freezing the encoder for the first epochs, a random 20% split for the validation dataset, an initial learning rates of 0.2 x 105 and a maximum of 50 epochs where the following epochs were used for experiments and analysis: “Aquilos 2” epoch 40, “Hydra” epoch 39 and “Aquilos 2 + Hydra” epoch 50. For the data from each microscope, validation and training datasets were determined only once, so that the same images were used for training and validation for both training and fine-tuning of the different models.

To allow users to train their own models using a command line interface, we implemented the model training and fine-tuning as described above in the open-source Python software “Adaptive Milling Model Training” (see Code Availability) together with instructions for installing and using the software, and how to annotate and prepare the data.

### Cell culture and vitrification

#### RPE-1 cells

Gold 200 mesh R2/2 UltrAuFoil grids (Quantifoil) were glow discharged and incubated for 1 hour with 150 µl polylysine (1 mg/ml) at room temperature. Grids were washed three times with Hanks’ Balanced Salt Solution (Gibco) and 0.18 × 10^6^ hTERT RPE-1 Tet-On:Plk4 GFP-hCent2 cells (Wang et al. 2011) were seeded on the grids with 1 ml DMEM/F12 medium (Thermo Fisher Scientific) with 10% FBS and cultured for 24 h. Cells were plunge-frozen using a Vitrobot Mark IV (Thermo Fisher Scientific) using blotting paper on the cell side of the grid and a polytetrafluoroethylene sheet on the backside of the grid. Just before blotting, 2 µl of DMEM/F12 medium (Thermo Fisher Scientific) with 10% FBS and 10% glycerol was applied to each grid. Vitrobot sample chamber was set to 37 °C at 80% relative humidity and blotting was performed with a blotforce of 10, blot time of 5 s and wait time of 2 s. After plunge-freezing, grids were directly clipped into Autogrids (Thermo Fisher Scientific).

#### Mouse embryonic stem cells

A HP1α-eGFP mESC stable cell line was generated via Lipofectamine 2000-mediated stable transfection of E14 mESCs with the pCAGGS-HP1α-eGFP-Puro plasmid, followed by puromycin selection, as previously described (Bulut-Karslioglu et al. 2014). These mESCs were cultured in petri dishes coated with 0.2% gelatin, in serum/LIF culture medium made up of 15% foetal bovine serum (Gibco), 1× NEAA (Gibco), 0.1 mM β-mercaptoethanol (Gibco), 1× Leukaemia inhibitory factor protein (Sigma-Aldrich), DMEM with Glutamax (Gibco). The cells were detached using TrypLE dissociation solution (Gibco) and resuspended into a homogeneous single cell suspension.

UltrAuFoil gold 200 mesh 2/2 EM grids (Quantifoil) were glow-discharged using a GloQube (Quorum) at 20 mA with negative polarity for 60 seconds and were briefly submerged in 70% ethanol (Fisher Scientific) for sterilisation and air dried for a few seconds. Sterilised grids were then pre-coated with 0.1 mg/ml poly-L-lysine (Sigma-Aldrich) prepared in foetal bovine serum (Gibco) at 37 °C for 30 minutes. 0.4 × 10^6^ cells were seeded onto the pre-coated EM grids ensuring even distribution and incubated at 37 °C for approximately 2 hours to allow cell attachment. EM grids with cells attached were vitrified in liquid ethane using a ThermoFisher Scientific Vitrobot Mark IV. Immediately prior to plunge freezing, the medium was replaced with cell culture medium containing 10% glycerol (Fisher Scientific), serving as a cryoprotectant to enhance vitrification efficiency. Grids were blotted using filter paper on both sides, with the blotting pads offset to predominantly blot from the rear.

#### Chlamydia infected HeLa cells

UltrAuFoil gold 200 mesh 2/2 EM grids (Quantifoil) were submerged in 70% ethanol for sterilisation and air dried for a few seconds. Sterilised grids were then transferred into a 35 mm Ibidi microscopy dish containing a shallow well filled with 1ml of FBS (Gibco) and 1% Gentamicin and left overnight in the incubator at 37 °C. The media in the dish was then replaced with cell culture medium containing DMEM with Glutamax and 10% FBS (Gibco) and 0.3 × 10^6^ HeLa cells were seeded onto the prepared EM grids. The cells were carefully pipetted directly onto the grids, ensuring even distribution across all grids. The grids were then incubated at 37 °C for 24 hours. After 24 h, HeLa cells were infected with C. trachomatis LGV2 serovar at an inclusion forming unit of 1, prepared by diluting bacterial stock in the infection medium. To synchronize infection, cells were centrifuged at 300 x g for 10 min at 4 °C and subsequently incubated at 37 °C with 5% CO2 for 80 min. After incubation, the infection medium was replaced with fresh medium, and cells were further incubated until plunge freezing at 24 and 48 hours post infection respectively. Just before plunge freezing, the medium in the dish was replaced with cell culture medium with 10% glycerol (Fisher Scientific), serving as a cryoprotectant. Grids were blotted for 8 s using a Leica GP2 (Leica Microsystems) operated at 37 °C and 70% humidity and immediately plunged into liquid ethane cooled by liquid nitrogen. Grids were clipped and stored under liquid nitrogen until imaging.

### (P)FIB lamella fabrication

#### Arctis

Some lamellae were prepared using an Arctis pFIB (Thermo Fisher Scientific) equipped with an ultra-high resolution (UHR) NICol non-immersion field-emission SEM column, using argon as the ion source. Once a suitable grid was loaded an overview image was taken using the Minimap tool in AutoLamella, the frontend GUI of fibsemOS. These SEM overview images were taken 90° relative to the grid at 2 kV, 0.1 nA with a 3 µs dwell time. Image size was 4096 x 4096 pixels with a field of view of 2800 x 2800 µm. The grid was then coated using a 120 s platinum deposition using the inbuilt microsputter. The FIB was adjusted to 12 kV and 0.15 µA for the 2-minute patterning of the platinum target. A 90 s GIS layer was then deposited before a second 120 s platinum layer. A second overview image was taken after coating, with the same imaging conditions to check for any grid damage from the GIS coating. The FIB beam alignments were run using the automatic alignment procedure from Thermo Fisher Scientific. These were run with the beam perpendicular to the back of the grid. A fiducial was first milled into a grid bar intersection to ensure there was sufficient contrast. The 30 kV currents in the range 20 pA to 2 nA were aligned. Beam shift alignments were also checked to ensure each current was centred on the same feature and all were close (*<* 5 µm) to coincident with the SEM. Suitable sites were then added as lamella sites in AutoLamella with a milling angle of 15°. For the infected HeLa cells, the same conditions were used, except that a milling angle of 13° was used.

Details of the milling steps are shown in Supplementary Table 2. All rough milling steps were carried out on all sites before moving onto fine milling and polishing. Note that the protocol was adjusted for the third conventional automation session as the grid was very well blotted and the 2 nA milling current was found to be overly destructive for the grid film. These steps were therefore reduced to 740 pA, which should not have a significant effect on the subsequent steps. For Adaptive Polishing SEM images were acquired with an SEM accelerating voltage of 2 kV and beam current of 25 pA using an Everhart–Thornley detector (ETD) with a dwell time of 50 ns, 16 frame integrations and a horizontal field width of 40 µm at an image resolution of 3072 x 2048 pixels. For Adaptive Polishing, the following settings were used: minimum GIS threshold of 250 nm, cracks/holes surface area threshold of 0.1 µm2, a maximum number of cycles of 60, SEM centring enabled, a maximum drift rate of 70 µm (effectively disabling it) and a minimum lamella area of 30 µm2. For Adaptive Polishing, regular automation used all milling steps except for the final polishing step, where the Adaptive Polishing step instead automatically adjusts the duration for each lamella site based on imaging feedback. For the conventional automation control, the same milling protocol was used, except that for the final polishing step the duration is the same for each site in an experiment. The time for the final polishing step for conventional automation was chosen to approximately match the median polishing time used by Adaptive Polishing session whilst also giving an even spread of polishing durations. For conventional automation, polishing times of 135, 270 and 405 seconds were used (Supplementary Table 2), which were chosen to roughly match the median Polishing times for the three Adaptive Polishing sessions (135, 255 and 375 seconds).

#### Helios Hydra

Some lamellae were prepared on a Helios Hydra G5 pFIB (Thermo Fisher Scientific) equipped with an Elstar SEM column, using argon as the ion source. The same workflow was applied with a few minor differences. The grids were loaded onto 27° pre-tilted shuttle. Grids were coated with 60 seconds of microsputter targeting a Pt stub at 12 kv 4 µA for 60 seconds, followed by 30 seconds of GIS coating and then another 60 seconds of microsputter. The FIB alignments were carried out at coincident height with the top of the grid perpendicular to the FIB, in a sacrificial area near the edge of the grid. A stage tilt of 4° (with stage rotation at -70°) was used to achieve the 15° milling angle. The ion column is the same between the Hydra and the Arctis, so the ion milling steps were the same, as shown in Supplementary Table 2. Adaptive Polishing settings were the same as with the Arctis.

#### Aquilos 2

Some lamellae were milled on an Aquilos 2 FIB (Thermo Fisher Scientific) at the electron Bio-Imaging Centre (eBIC). The Aquilos 2 is equipped with the same NICol UHR non-immersion field emission SEM column as the Arctis. The same workflow was applied again with a few minor differences. The grids were loaded onto a 26° shuttle. Sputter coating was performed at 1 kV, 30 mA, 60 s, 0.1 mbar argon, followed by 30 or 50 seconds of GIS coating and then another sputter layer at the same settings. The milling angle used was 12°. These settings were in line with those usually used by eBIC users. The Aquilos has a gallium ion column, so the milling currents used were slightly different (Relief cuts: 1 nA, Fiducial: 100 pA, Rough 1: 1 nA, Rough 2: 1 nA, Fine 1: 500 pA, Fine 2: 50pA, Fine 3: 30 pA and Adaptive Polishing: 30 pA). Adaptive Polishing settings were the same as with the Arctis. To collect more training data of cracks and holes, for part of the lamellae prepared on the Aquilos 2 the GIS thickness threshold was lowered to 0 and the hole area threshold value increased to 2.1 µm2, to effectively only stop polishing once very large cracks/holes have formed. In the subsequent acquisition session, reasonable thresholds were used in a first Adaptive Polishing step, followed by a second Adaptive Polishing step with the above values to destroy the lamellae.

### Cryo-electron tomography

For the comparison of Adaptive Polishing with conventional automation, data was collected on a Titan Krios G4 (Thermo Fisher Scientific) equipped with a Falcon 4i camera and Selectris X energy filter. Using Tomo5 software (Thermo Fischer Scientific), low-dose overviews were collected at 15° to correct for the lamella pre-tilt with a pixel size of 56.2 Å (magnification of 2250 x) with a 200 µm defocus. To minimise sampling biases during cryo-ET data acquisition and to account for the lamella wedge-like geometry(Schaffer et al. 2017; Berger et al. 2023, 2025; Tuijtel et al. 2024), 12 tilt series were acquired per lamella in a fixed pattern irrespective of local sample thickness or biological features. For each lamella, 12 dose-symmetric(Hagen et al. 2017) tilt series were acquired in parallel using beam shift(Khavnekar et al. 2023; Eisenstein et al. 2023), where the centre of 4 tilt series were placed approximately 1 µm away from the GIS layer, 4 tilt series in the approximate middle of the lamella, and the remaining 4 approximately 1 µm away from the grid foil (Supplementary Fig. 5). 360 and 432 tilt series were collected in total over 3 independent sessions on lamellae prepared with Adaptive Polishing and conventional automation respectively. Tilt series were collected in electron counting mode in EER format at a calibrated pixel size of 1.90 Å (magnification of 64,000 x), with a starting tilt range of ±51° with 3° increments starting at 15°, a dose per tilt of 3.63 e^-^/Å^2^ (35 tilt images with 127 e^-^/Å^2^ total dose), with a defocus range between 3 and 5 µm in 0.5 µm steps.

For the infected HeLa cells, a total of 714 tilt series were collected on the same instrument using the same pixel size and defocus range, but positions were chosen based on biological relevance and lamella quality rather than in a fixed pattern. For the grids frozen 24 h post infection, tilt series were collected with a tilt range of ±51° with 3° increments starting at -13°, a dose per tilt of 4 e^-^/Å^2^ (35 tilt images with 140 e^-^/Å^2^ total dose) was applied. Tilt series of grids frozen 48 h post infection were collected with the same conditions except that a tilt range of ±39° with 3° increments starting at -13° was used and a dose per tilt of 3 e^-^/Å^2^ (27 tilt images with 81 e^-^/Å^2^ total dose) was applied.

### Tomography data processing

For the data from the comparison of Adaptive Polishing to conventional automation, tilt images of low quality (e.g. low contrast or objects moving into the field of view) were excluded from tomography processing using a custom script and WarpTools (https://github.com/warpem/warp) (Tegunov et al. 2021) was used for gain correction, CTF estimation, and motion correction with a frame group size of 9. Tilt series were aligned using AreTomo (Zheng et al. 2022) (version 1.2.5) and tomograms were reconstructed with a binning factor of 8 using WarpTools. An IsoNet (Liu et al. 2022) model was trained on a total of 3216 subvolumes extracted from 804 unmasked tomograms using a cube size of 32 and a crop size of 128 and applied to the tomograms. These tomograms were used for measuring the local lamella thickness using the measure tool in IMOD (Mastronarde and Held 2017). For the infected HeLa cells data, the data was processed similarly, except that an IsoNet model was trained from using a total of 1000 subvolumes only from 10 tomograms.

### Milling success rate and lamella quality evaluation

Lamella milling outcomes were assessed using TEM overview images. Lamella length, crack length, lamella surface area, and hole area were manually measured in FIJI (Schindelin et al. 2012). Each lamella site milled was classified into one of four categories: Excluded, Failed, Partial or Success.

Lamellae were classified as excluded when failure occurred prior to the polishing step, such as breakage during earlier milling stages, as these events were typically associated with incorrect lamella positioning and therefore attributed to user or sample preparation errors. Specific cases include: 1) Lamellae that were not successfully thinned due to suboptimal user input such as positioning too close to grid bars or not correctly set at the coincident height, 2) lamellae with sufficient remaining back-ice to partially obscure the lamella in TEM when viewed perpendicular to the electron beam, and 3) lamellae that were out of the stage limits of the TEM and so tilt series could not be collected.

Lamellae were classified as failed if they were destroyed during the polishing step or during transfer to the TEM, contained a double lamella, exhibited severe damage, or remained visibly too thick due to incorrect decisions during the Adaptive Polishing step. Severe damage was defined as either a full-length crack extending along the lamella (resulting from milling or transfer) or holes occupying more than 20% of the total lamella area.

Lamellae were classified as partial successes when they exhibited cracks extending between 50% and 100% of the lamella length (excluding full-length cracks) or holes occupying 5–20% of the lamella area. Successful lamellae were defined as those that remained fully intact or contained only minor defects, including cracks shorter than 50% of the lamella length or holes occupying less than 5% of the lamella area. Tilt series were collected where possible, regardless of the status of the lamella.

Lamella thickness was measured using the line measurement tool in IMOD (Mastronarde and Held 2017) by flipping the tomogram, guided by the XYZ viewer, to get the edge on view. The line measurement was taken at the approximate centre of the tomogram. This was done for all acquired tomograms, unless the quality was too low to reliably measure the local lamella thickness.

## Movies

### Movie 1 and 2

Overview of acquired images, segmentations and decisions made by Adaptive Polishing on the microscope, for the two examples shown in Fig. 1. The milling time shown at the top is per milling pattern (30 seconds milling time between each overview) and the final overview is where Adaptive Polishing was stopped, because one of the threshold criteria was reached (below 250 nm GIS thickness or cracks/holes area above 0.10 µm2). The left panels show the acquired SEM and FIB images. The middle panel shows the segmentation (lamella: green, GIS: orange, cracks/holes: red, vacuum: purple, background: blue). The top-right panel shows the segmentation after post-processing, with vertical purple lines showing the lamella width used for measuring the minimum GIS thickness. Bottom right plot shows the local GIS thickness, with the dashed orange line showing the GIS threshold at which Adaptive Polishing stops.

### Movie 3-6

Tomograms of the slices shown in Fig. 2e-h respectively. Scale bar: 100 nm.

#### Movie 7-8

Overview of acquired images, segmentations and decisions made by Adaptive Polishing with the “All” model on a Helios Hydra (Movie 7) and the “Aquilos 2” model on an Aquilos 2. “All” and “Aquilos 2”.

#### Movie 9-10

SEM images of the lamellae shown in Figure 4c and d segmented with the 5 models indicated at the top. No post-processing was applied to these segmentations.

## Statistics and Reproducibility

Unless otherwise stated, data are presented as mean±SD. Statistical significance was assessed using two-sided Welch’s t-tests, which do not assume equal variances, for which P-values *<* 0.05 were considered statistically significant. Exact sample sizes (n) are indicated in the corresponding figure legends or the Results. No statistical methods were used to predetermine sample sizes. Experiments comparing Adaptive Polishing with conventional automation were performed in three independent technical replicate milling sessions. For lamella thickness measurements, n refers to individual tomograms. For milling success rates, n refers to individual lamellae.

## Data availability

The lamella SEM images and segmentations together with all the pre-trained models are available on Zenodo (DOI: 10.5281/zenodo.21804785) licenced under CC BY 4.0. The cryo-ET data acquired for comparing Adaptive Polishing and conventional automated milling performance will be deposited on EMPIAR licenced under CC0.

## Code availability

The Adaptive Milling package including the Adaptive Polishing plugin licenced under Apache 2.0 with the Commons Clause is available on GitHub (https://github.com/rosalindfranklininstitute/adaptive_milling) together with user guides. The “Adaptive Milling Model Training” packaged licensed under Apache 2.0 with the Commons Clause is available on GitHub (https://github.com/rosalindfranklininstitute/AM-modeltraining).

## Acknowledgements

We thank Jennifer Wang, Garrison Kent Bus and Tim Stearns for providing the hTERT RPE-1 Tet-On:Plk4 GFP-hCent2 cell line and providing cell culture protocols. We thank Michelle Percharde and Kamila Musialik for providing the HP1α-eGFP mESC stable cell line and cell culture protocols. We thank Matthew Case for TEM microscope support, Matt Spink for FIB/SEM support, and Avery Pennington for discussions to optimise segmentation model training. We thank all early testers of Adaptive Polishing: Kathryn Smyth, Becky Csondor, William Bowles, Angharad Smith, Siva Ramadurai, Victoria Garcia Giner, Miron Leanca, and Jianguo Zhang.

This work was supported by the Wellcome Trust through the Electrifying Life Science Project (220526/Z/20/Z). The authors acknowledge the support and the use of resources of Instruct-ERIC through the TechDev pilot scheme APPID 4119. We acknowledge Diamond Light Source Ltd. for access and support of the cryo-EM facilities at eBIC, proposals nr30048, cm44126, and cm40597. Funding was provided to P.C. from the Chan Zuckerberg Initiative, grant number 2025-366327. The Rosalind Franklin Institute is funded by the UK Research and Innovation, Engineering and Physical Sciences Research Council.

## Author contributions

T.G. and CB conceived the study. N.G., R.T.R and C.B. prepared cryo samples. T.G., R.T.R and C.B. prepared (plasma) FIB lamellae. R.T.R and C.B. collected and processed cryo-ET data. C.B. segmented SEM data. T.F. L.P. and C.B. trained machine-learning segmentation models. T.F., E.H., P.C. and L.P. wrote the software. T.G., T.F. and C.B. analysed the data, wrote the manuscript and prepared figures. M.D., M.B., M.D. and C.B. supervised the project. All authors reviewed the manuscript and the data.

