## Supplementary Data for "Adaptive Milling: imaging feedback-driven automated fabrication of consistently thin cryo-lamellae"

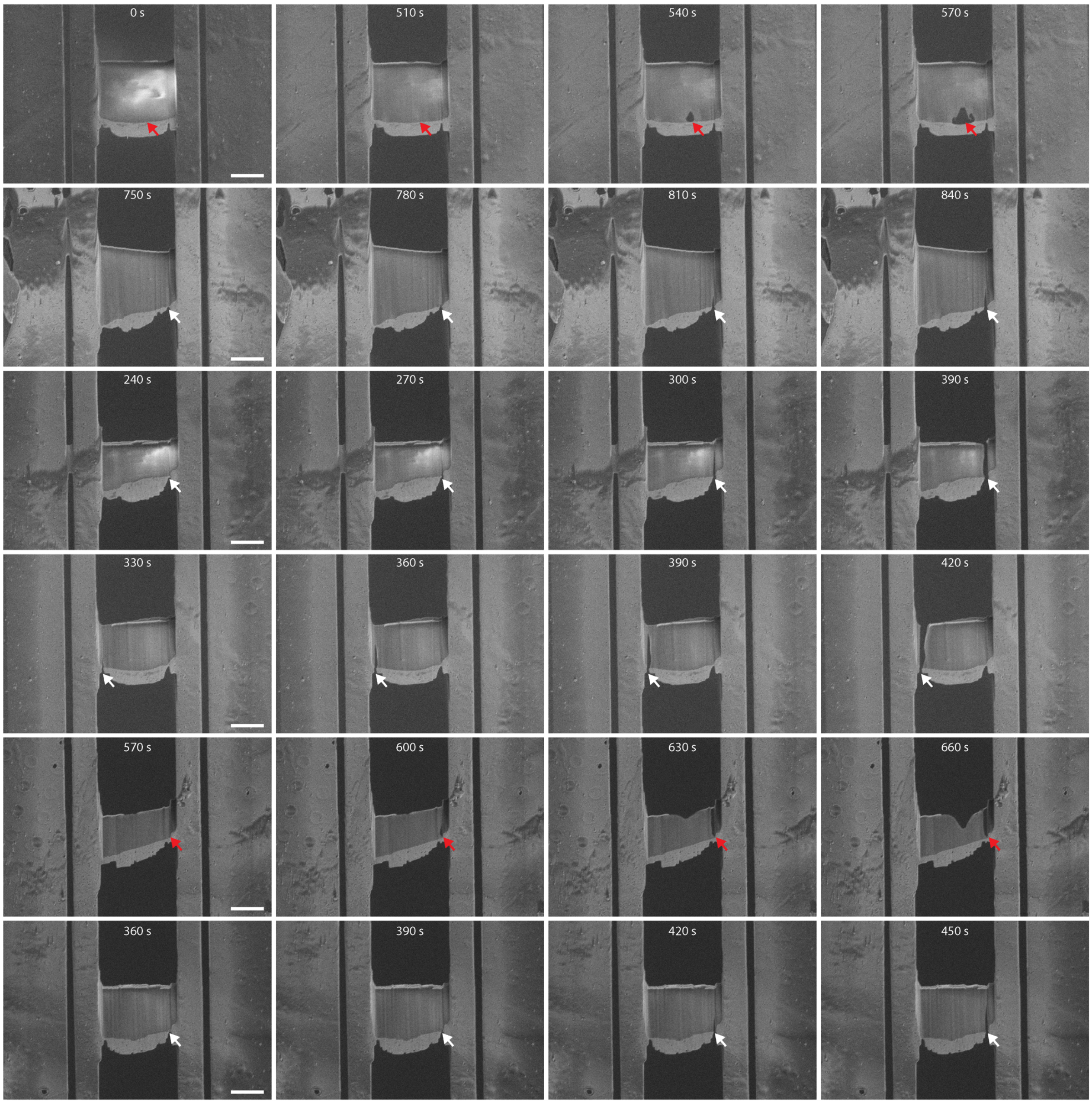


***Supplementary Figure 1. Examples of crack and hole formation during polishing.*** *SEM images acquired during polishing of 6 different lamellae. White arrows indicate positions where cracks are rapidly formed when polishing is resumed when the GIS has locally been removed. In some more rare cases, holes form in the lamella when there is still GIS directly in front of it (red arrows). The total polishing time in seconds for each SEM image is displayed at the top. Scale bars: 5 µm.*

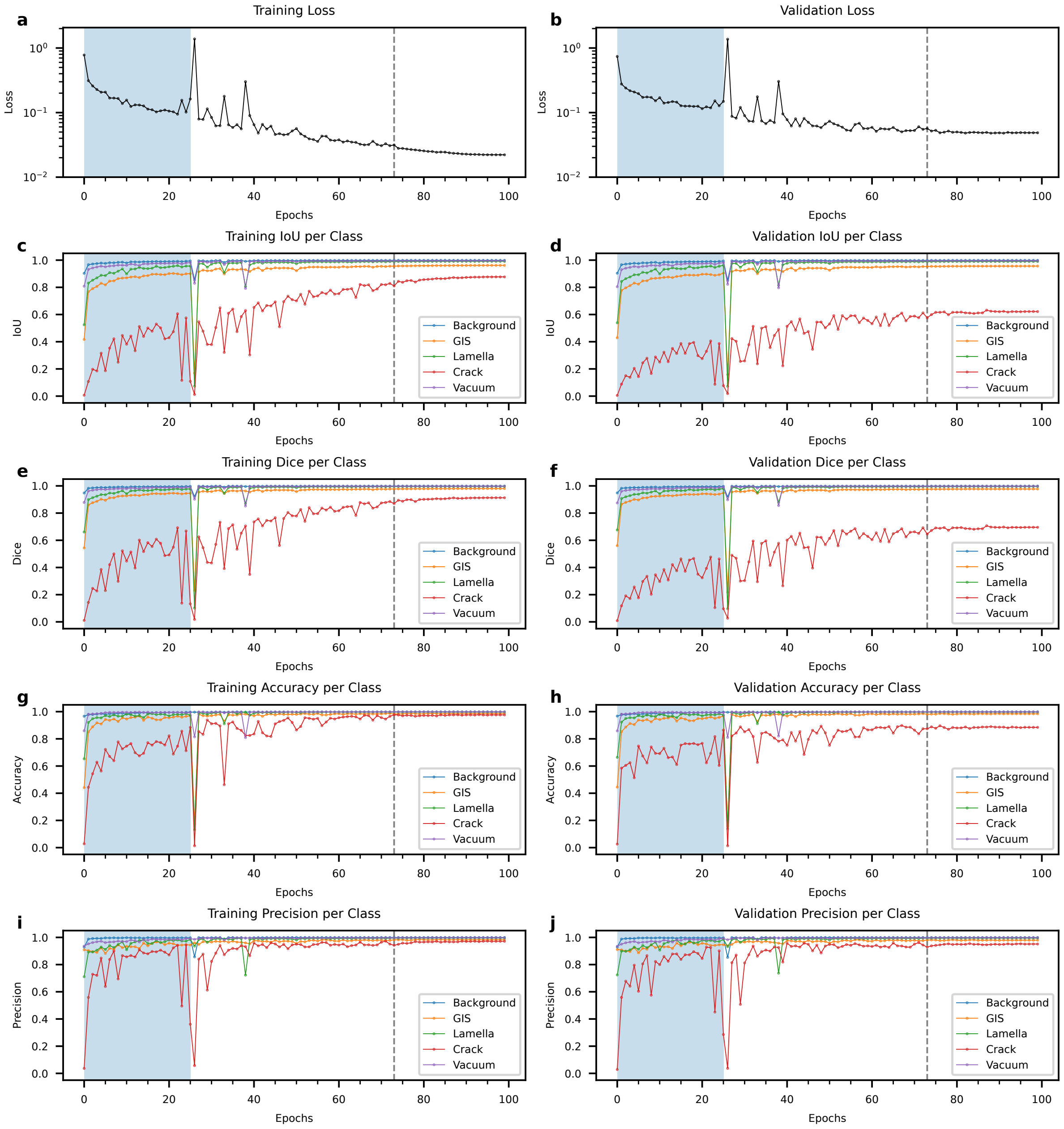


***Supplementary Figure 2. Model performance metrics during training.*** *Training and validation metrics for the model trained on 2,873 SEM images of lamellae labelled with background, the GIS layer, the lamella surface area, cracks/holes and vacuum. Plots display training (left) and validation (right) metrics over the different epochs for each of the 5 labelling classes. The encoder was frozen for the first 25 epochs (blue background). Epoch 73 (vertical grey dashed line) was the model used for subsequent microscope tests. Plotted metrics are: (a,b) Compound loss function of weighted cross-entropy (50%) and weighted multiclass Dice loss (50%). (c,d) intersection over union (IOU) scores, (e,f) Dice coefficient, (g,h) accuracy and (i,j) precision.*

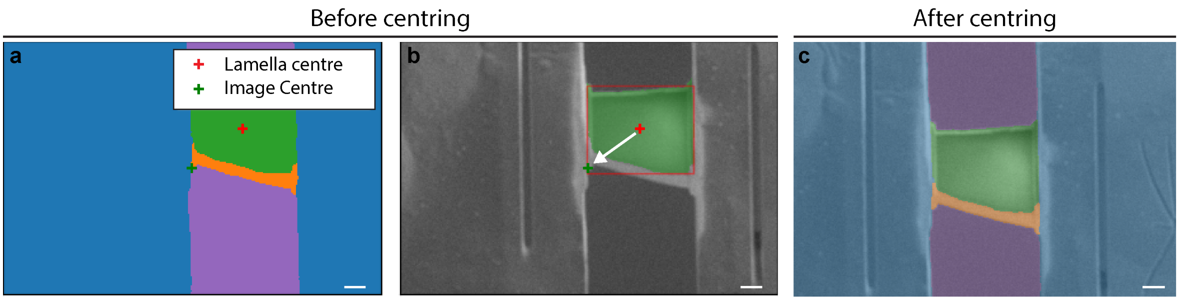


***Supplementary Figure 3. Routine for centring lamellae on the field of view of the SEM.*** *(a,b) An SEM image is acquired and segmented using the machine-learning model. This is used to determine the centre of the lamella (red +). An SEM beam shift (white arrow) is than applied to move the centre of the lamella towards the centre of the field of view (green +). (c) overlay of SEM image with segmentation, acquired after applying this beam shift. Scalebars: 2 µm.*

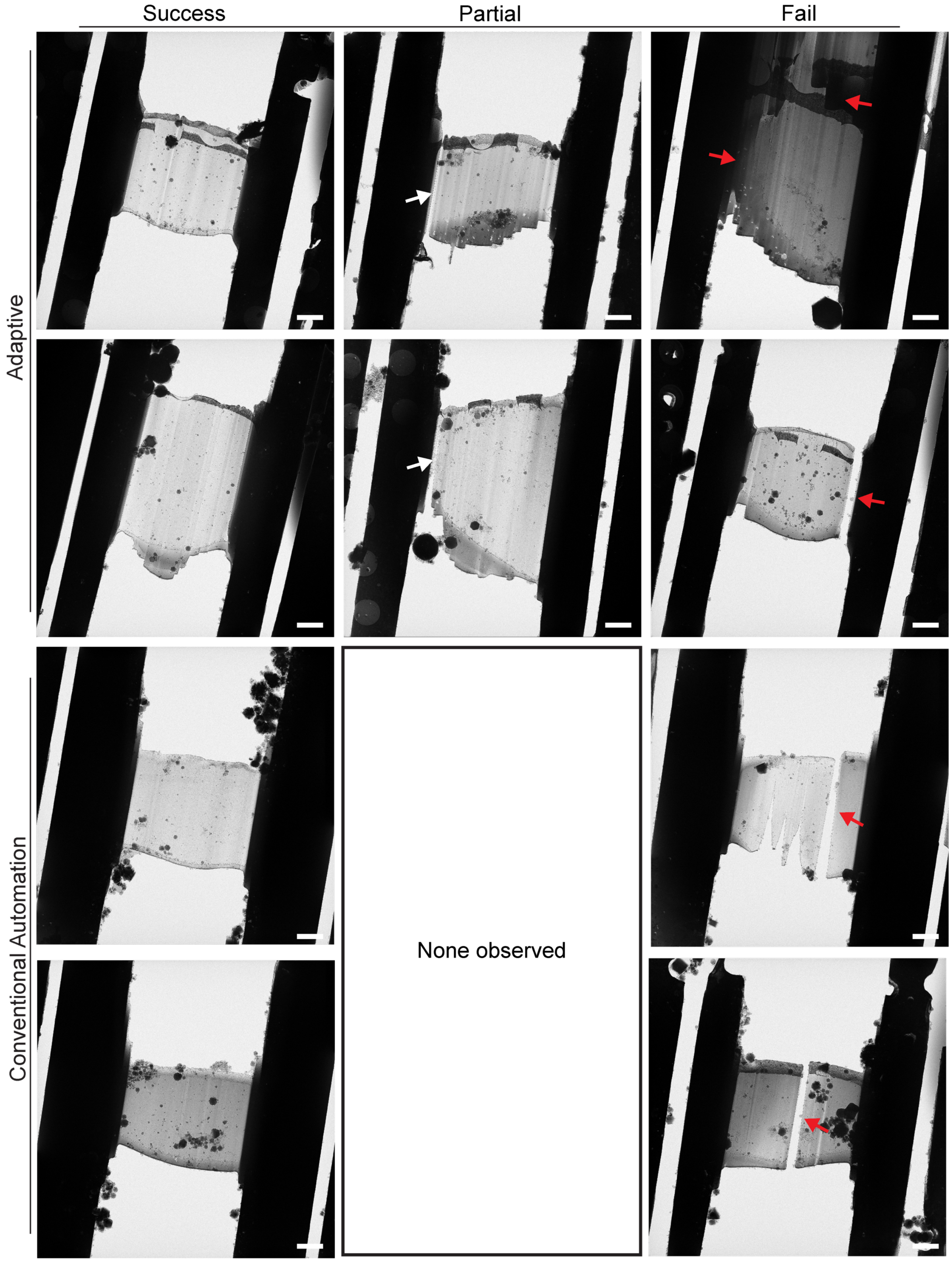


***Supplementary Figure 4. Examples of lamella and lamella classification.*** *TEM images acquired before tilt series acquisition. White arrows indicate positions where cracks more than half the length of the lamella have formed during polishing leading to classification of 'partial success'. Red arrows indicate full length cracks or issues with double lamella. There were no instances of 'partial success' during the control experiments with conventional automation. Scale bars: 2 µm.*

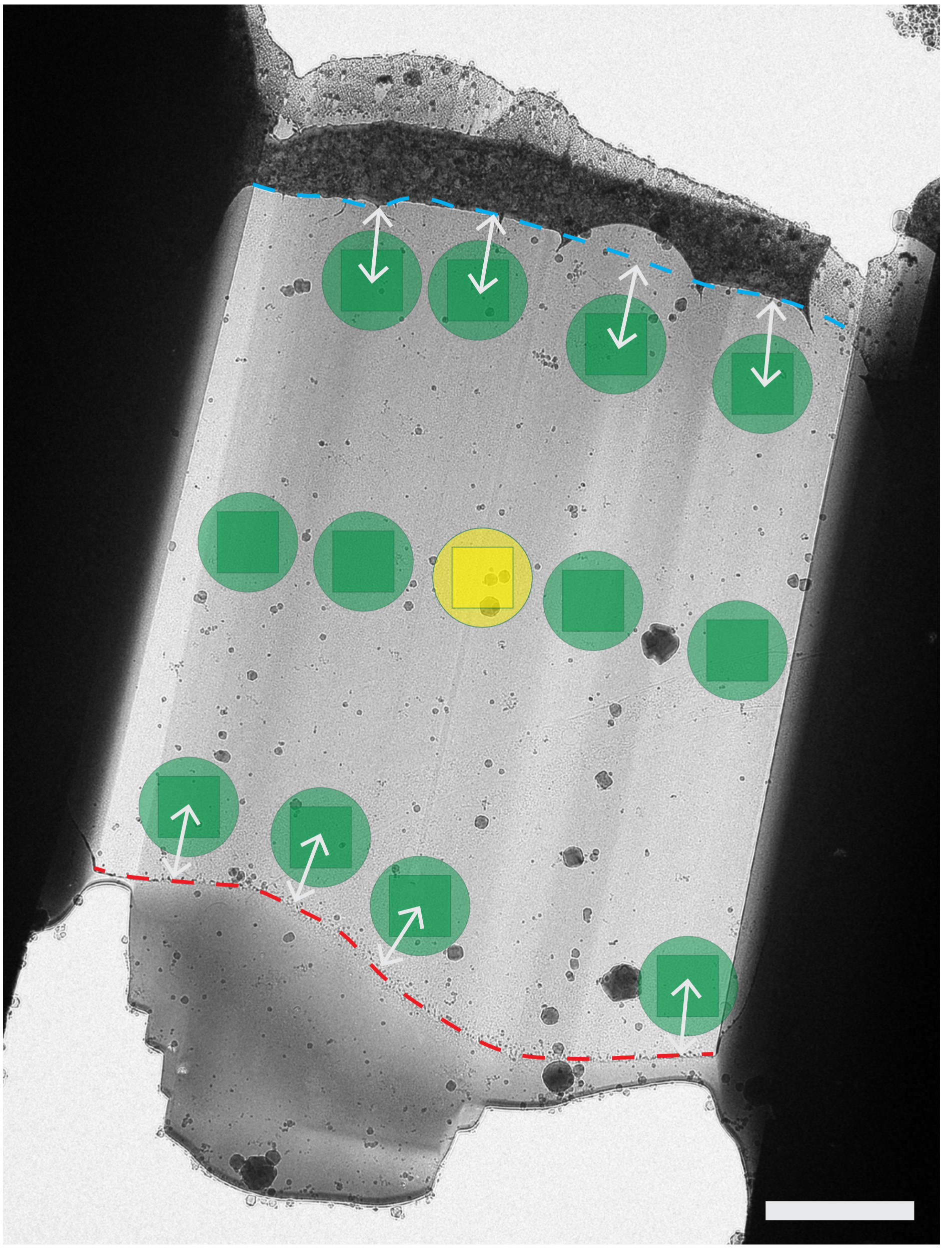

***Supplementary Figure 5. Example TEM Lamella overview showing pattern in which tilt-series were acquired.*** *A total of 12 tilt-series were collected per lamella, placed in 3 rows of 4 tilt-series each. Tilt-series field of view is indicated as green squares, with the beam diameter as a green circle, and the focus/tracking area in yellow. One row of tilt-series was placed at the approximate centre of the lamella, with one row placed approximately 1 µm away (white double-headed arrows) from the GIS-lamella interface (red dashed line), and the final row approximately 1 µm away from the edge of the grid foil (blue dashed line). Tilt-series were placed to avoid large ice contaminations, but thicker lamella regions were not avoided, and thinner regions not favoured. Scale bars: 2 µm.*

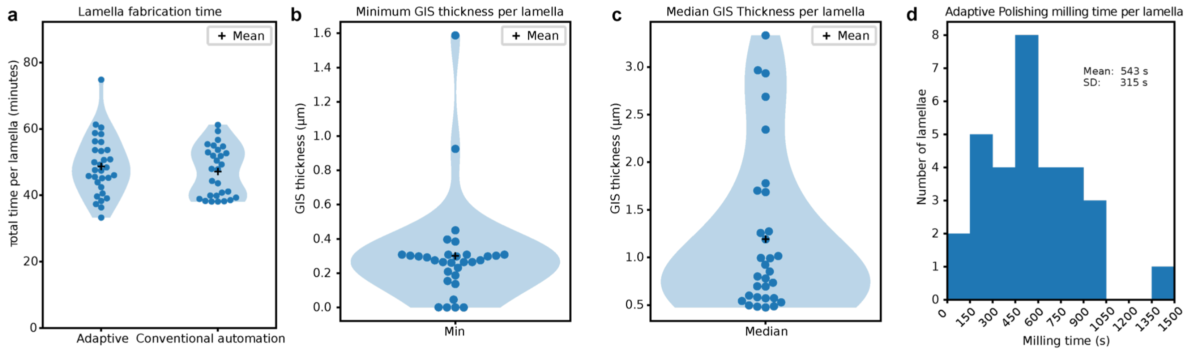


**Supplementary Figure 6. Per lamella statistics for Adaptive Polishing.** *(a) Total duration of automated lamella fabrication for both Adaptive Polishing and conventional automation. For Adaptive Polishing, the mean±SD duration for 31 lamellae was 15m15s±8m09s compared to 9m09s±3m04s for the 28 lamellae prepared with conventional automation. (b) Distribution of the minimum GIS thickness after Adaptive Polishing. (c) distribution of the median GIS thickness after Adaptive Polishing. (d) Histogram of the time spent polishing every lamella during the Adaptive Polishing step (excluding imaging and other overhead).*

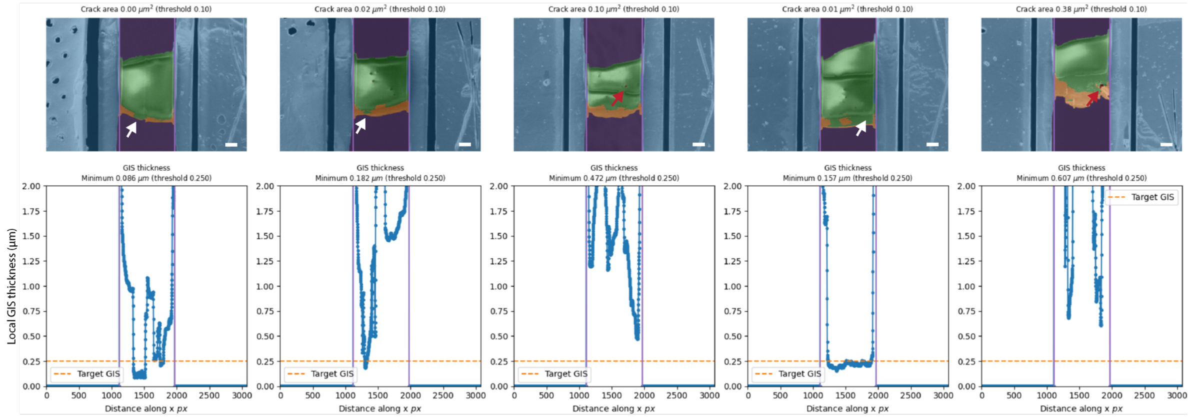


***Supplementary Figure 7. Examples of incorrect decision-making by Adaptive Polishing on the Helios Hydra due to the Arctis model.*** *Example lamellae which were stopped because of segmentations in the GIS-lamella boundary (white arrows) or false-positive holes (red arrows). Scalebars: 2 µm.*

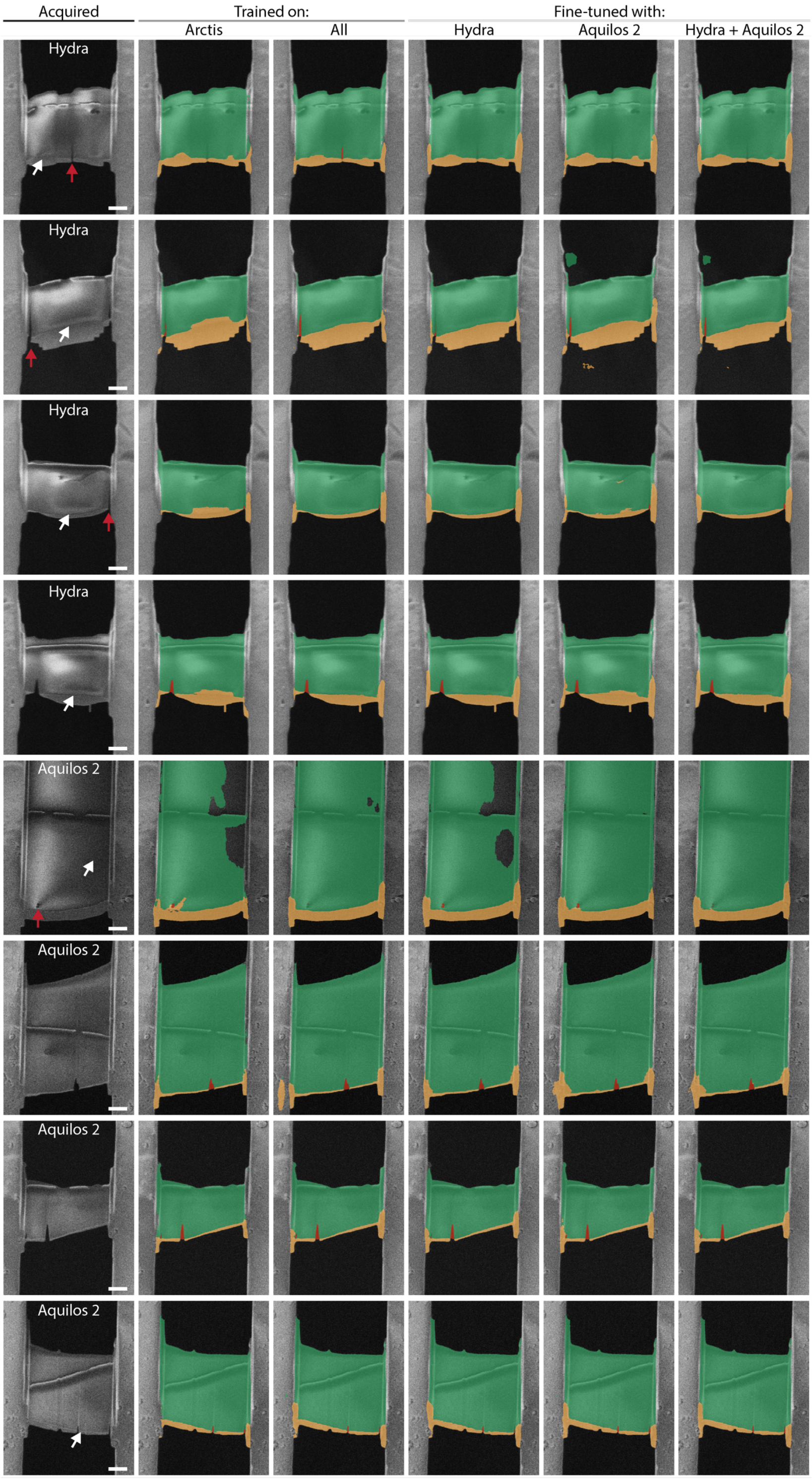


***Supplementary Figure 8. Model performance on other microscopes after training or fine-tuning.*** *SEM images acquired on Helios Hydra and Aquilos 2 FIB/SEM microscope, with the predictions (without post-processing) from the original model trained on the Arctis, the model trained with data from all three microscopes, and three models fine-tuned with data from the Aquilos 2, Helios Hydra, or both. All of the lamellae shown in this figure were created after model training and fine-tuning. White and red arrows indicate areas where notable improvements in segmentations with some of the models can be observed. Scalebars: 2 µm.*

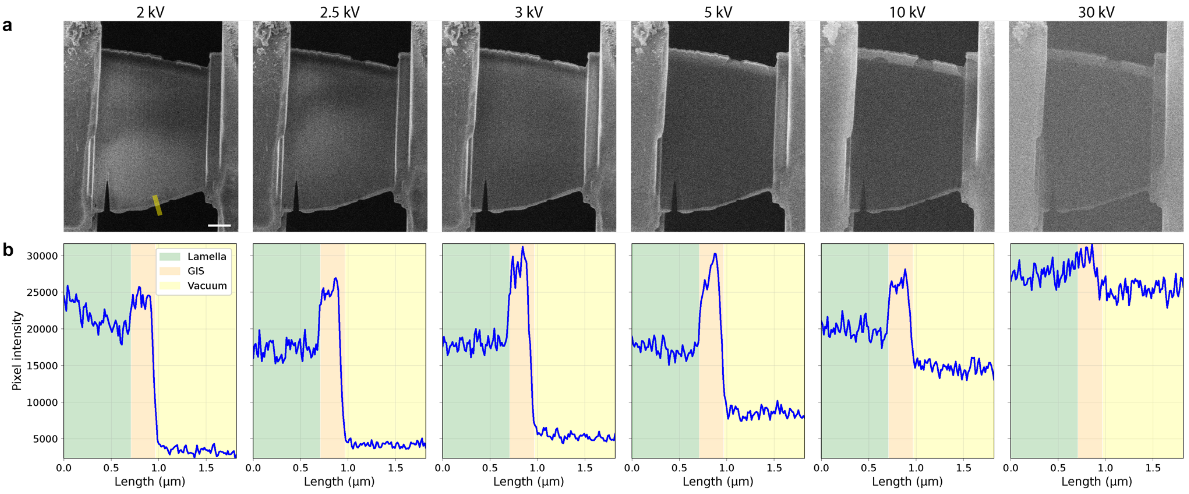


c

| Ratio | 2kV | 2.5kV | 3kV | 5kV | 10kV | 30kV |
|---|---|---|---|---|---|---|
| GIS/lamella | 1.041 | 1.262 | 1.359 | 1.468 | 1.256 | 1.041 |
| lamella/vacuum | 6.211 | 4.115 | 3.372 | 2.011 | 1.347 | 1.086 |
| GIS/vacuum | 6.467 | 5.195 | 4.582 | 2.953 | 1.692 | 1.131 |

***Supplementary Figure 9. Effects of the acceleration voltage on contrast between the GIS layer, lamella surface and vacuum on an Aquilos 2.*** *(a) SEM images acquired at different acceleration voltages on an Aquilos 2. Yellow line indicates where a 50-pixel wide line profile was measured, after image alignment with SIFT. Substantial differences in contrast can be observed in the lamella surface area, GIS layer and vacuum as a function of the acceleration voltage. Scalebar 2 µm. (b) Line intensity profiles of the pixel intensity measured along the line in panel a. Colours indicate the range corresponding to the lamella, GIS and vacuum areas. (c) colour-coded table showing mean pixel intensity ratios of the coloured areas in panel b. The colour-coded values of each of the three ratios is scaled separately. Contrast between the lamella and GIS with the vacuum is highest at lower acceleration voltages, whereas the highest contrast is observed between the GIS and the lamella at around 5 kV.*

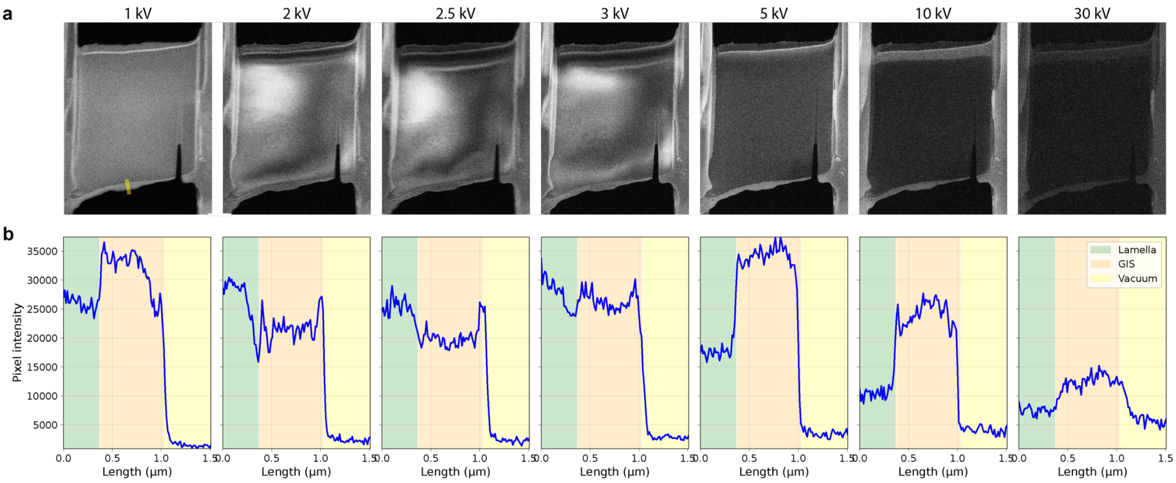


c

| Ratio | 1kV | 2kV | 2.5kV | 3kV | 5kV | 10kV | 30kV |
|---|---|---|---|---|---|---|---|
| GIS/lamella | 1.206 | 0.825 | 0.799 | 0.951 | 1.798 | 2.120 | 1.646 |
| lamella/vacuum | 12.966 | 9.252 | 5.618 | 7.589 | 5.243 | 2.687 | 1.106 |
| GIS/vacuum | 15.633 | 7.636 | 4.488 | 7.219 | 9.426 | 5.697 | 1.820 |

***Supplementary Figure 9. Effects of the acceleration voltage on contrast between the GIS layer, lamella surface and vacuum on Helios Hydra.*** *(a) SEM images acquired at different acceleration voltages on a Helios Hydra. Yellow line indicates where a 50-pixel wide line profile was measured, after image alignment with SIFT. Substantial differences in contrast can be observed in the lamella surface area, GIS layer and vacuum as a function of the acceleration voltage. Scalebar 2 µm. (b) Line intensity profiles of the pixel intensity measured along the line in panel a. Colours indicate the range corresponding to the lamella, GIS and vacuum areas. (c) colour-coded table showing mean pixel intensity ratios of the coloured areas in panel b. The colour-coded values of each of the three ratios is scaled separately. Contrast between the lamella and GIS with the vacuum is highest at 5 and 10 kV, with the signal intensity of the GIS being lower than the lamella between 2 and 3 kV.*

**Supplementary table 1. Overview of reported lamella thicknesses in the literature.**

| Reference | Milling method | Sample | Thickness | notes |
|---|---|---|---|---|
| Zachs et al. 2020[1] | Automated and manual gallium FIB | Yeast and cyanobacteria | Mean: 243 nm *(automated)*<br>Mean: 258 nm *(manual)* | Thickness measured using cryo-ET. |
| Berger et al. 2021[2] | Manual with gallium FIB | Primary human myeloid cells infected with *yersinia enterocolitica* | Mean±SD: 186±58 nm | Thickness measured manually in IMOD in all 63 tomograms used for STA (>150 collected). |
| Klumpe et al. 2021[3] | Automated with a gallium FIB | Mammalian cells, *Emiliania huxleyi*, yeast and *Chlamydomonas reinhardtii* | mean:<br>~245 & ~265 nm *(HeLa cells)*<br>~340 & ~250 nm (*Sum159 cells)*<br>~340 and ~330 nm (*E.huxleyiI)*<br>~340 & ~350 nm (*C. reinhardtii*) | Thickness measured using cryo-ET. |
| Li et al. 2023[4] | Manual with gallium FIB | HeLa cells | Mean±SD: 169±43 nm | Thickness measured on 46/75 tomograms. |
| Berger et al. 2023[5] | Automated with argon pFIB | HeLa cells | Mean±SD: 250±70 nm<br>Mean±SD: 227±62 nm<br>Mean±SD: 245±76 nm | Thickness measured with script from manually annotated surface models on all acquired tilt-series (180, 30 and 47 respectively). |
| Yang et al. 2023[6] | Gallium FIB at 30 and 8 kV | Parvovirus and yeast | Range: 76 – 296 nm (virus *30 kV*)<br>Range: 60 - 266 nm (virus *8 kV)*<br>Range: 78 - 400 nm (*yeast 8kV)* | Thickness measured on 216 and 110 parvovirus tomograms and thickness estimated on 183 micrographs of yeast using the Beer-Lambert law. |
| Tuijtels et al. 2024[7] | Automated with manual polishing<br>With a gallium FIB | *Dictyostelium discoideum* | Range: 43 - 255 nm<br>Mean: 155 nm | Thickness measured manually in IMOD on 261/343 of the acquired tilt-series. |
| Schiøtz et al. 2024[8] | Manual gallium FIB milling of Serial lift-out chunks | *Caenorhabditis elegans* L1 larvae | Mean±SD: 255±126 nm (*single-side attachment*)<br>303 ± 40 nm and 252 ± 36 nm (*double-sided*) | Thickness measured on all 56 tomograms for single-side attached lift-out chunks, and two technical replicates of double-sided attachment with a random selection of 132/1012 tomograms, and 90/90 respectively. |
| Nguyen et al. 2024[9] | SOLIST lift-out chunks with gallium FIB milling | Yeast | Mean±SD: 184±9.3 nm (*lift-out*)<br>Mean±SD: 187±31 nm (*SOLIST*) | Thickness was measured in at least two tomograms per lamella using IMOD, at three positions per tomogram for 6 lift-out and 32 SOLIST lamellae. |
| Berger et al. 2025[10] | Automated with xenon pFIB, with some manual supervision during polishing | High-pressure frozen *Escherichia coli* and yeast (waffle-style) | Mean±SD: 201±34 nm (*E. coli)*<br>Mean±SD: 144±45 nm (*yeast)*<br>Mean±SD: 209±42 nm (*yeast)* | Thickness measured with script from manually annotated surface models. 167/286, 216/369 and 96/160 tomograms respectively. |
| Kelley et al. 2026[11] | Automated with xenon and argon pFIB | *Chlamydomonas reinhardtii* | Median: 157 nm | Thickness measured with script for 1829/2991 tomograms, with further 10% outliers removed. |
| This study | Automated Adaptive Polishing with argon pFIB | Mouse embryonic stem cells | Mean±SD: 158±42 nm | Thickness measured manually with IMOD for 328/360 tomograms. Tomograms were acquired in fixed pattern to avoid selection bias. |
| This study | Automated with argon pFIB | Mouse embryonic stem cells | Mean±SD: 235±78 nm | Thickness measured manually with IMOD for 271/432 tomograms. Tomograms were acquired in fixed pattern to avoid selection bias. |
| This study | Automated Adaptive Polishing with argon pFIB | HeLa cells infected with *C. trachomatis* LGV2 serovar | Mean±SD: 150±39 nm | Thickness measured manually with IMOD for 674/714 tomograms. |

*Overview of studies reporting the thicknesses of cryo-lamellae, and which milling methods and samples were used.*

**Supplementary Table 1. Milling parameters used for assessing Adaptive Polishing performance on the Arctis.**

| Milling step | Current | Pattern dimensions (h x w) (µm) | Milling depth (µm) | Distance between milling patterns | Milling time |
|---|---|---|---|---|---|
| Relief cuts | 2 nA / 740 pA | 18 x 1 / 10 x 0.8 | 2.25 OR 1.5 | NA | 0m 50s / 0m 40s |
| Fiducial | 60 pA | NA | 0.50 | NA | 1m 34s |
| Rough 1 | 2 nA / 740 pA | 6 x 11 | 2.50 OR 2.0 | 6 µm | 5m 06s / 11m 51s |
| Rough 2 | 740 pA | 3 x 11 | 2.40 | 3 µm | 7m 38s |
| Fine 1 | 200 pA | 1.25 x 11 | 1.40 | 1.3 µm | 6m 04s |
| Fine 2 | 60 pA | 0.5 x 10 | 0.65 | 600 nm | 3m 26s |
| Fine 3 | 20 pA | 0.25 x 10 | 0.40 | 400 nm | 4m 14s |
| Adaptive Polishing<br>OR<br>Polish | 20 pA<br><br>20 pA | 0.35 x 10<br><br>0.35 x 10 | NA<br><br>NA | 200 nm<br><br>200 nm | Various<br><br>2m 15s / 4m 30s / 6m 45s |

*The polish step was used instead of the Adaptive Polishing step for conventional automation. All milling was done with argon as the ion source with an acceleration voltage of 30 kV. Some changes were made for one of the conventional automation experiments as the grid was very well blotted and reducing the current from 2 nA to 740 pA was found necessary to avoid destroying the grid film. The conventional automation experiment used the polish step, whilst the adaptive run used the Adaptive Polishing step instead.*
